# Dynamic Causal Modeling in Probabilistic Programming Languages

**DOI:** 10.1101/2024.11.06.622230

**Authors:** Nina Baldy, Marmaduke Woodman, Viktor Jirsa, Meysam Hashemi

## Abstract

Understanding the intricate dynamics of brain activities necessitates models that incorporate causality and nonlinearity. Dynamic Causal Modelling (DCM) presents a statistical framework that embraces causal relationships among brain regions and their responses to experimental manipulations, such as stimulation. In this study, we perform Bayesian inference on a neurobiologically plausible generative model that simulates event-related potentials observed in magneto/encephalography data. This translates into probabilistic inference of latent and observed states of a system driven by input stimuli, described by a set of nonlinear ordinary differential equations (ODEs) and potentially correlated parameters. We provide a guideline for reliable inference in the presence of multimodality, which arises from parameter degeneracy, ultimately enhancing the predictive accuracy of neural dynamics. Solutions include optimizing the hyperparameters, leveraging initialization with prior information, and employing weighted stacking based on predictive accuracy. Moreover, we implement the inference and conduct comprehensive model comparison in several probabilistic programming languages to streamline the process and benchmark their efficiency. Our investigation shows that model inversion in DCM extends beyond variational approximation frameworks, demonstrating the effectiveness of gradient-based Markov Chain Monte Carlo methods. We illustrate the accuracy and efficiency of posterior estimation using a self-tuning variant of Hamiltonian Monte Carlo and the automatic Laplace approximation, effectively addressing parameter degeneracy challenges. This technical endeavor holds the potential to advance the inversion of state-space ODE models, and contribute to neuroscience research and applications in neuroimaging through automatic DCM.

## 1. Introduction

Dynamic Causal Modelling (DCM; Friston et al. (2003); Razi et al. (2015); Frässle et al. (2017)) has become a cornerstone methodology in neuroimaging for understanding and interpreting complex brain dynamics. Central to DCM is Bayesian model inversion on a set of ordinary differential equations (ODEs), which aims to infer the posterior distribution of lumped parameters given the prior and observed data (van de Schoot et al., 2021; Gelman et al., 2014a; Box and Tiao, 2011). This process is challenged by the complexity and high-dimensionality of the models, often rendering evaluation of the posterior density computationally intractable (Smith, 1991). To address this issue, one common approach is to use a fixed-form probability distribution to approximate the posterior density, which relaxes the challenging computation of high-dimensional integrals into an optimization problem. Traditionally, variational inference (VI; Jordan et al. (1999); Blei et al. (2017)) has been the preferred approach for this task, due to its computational efficiency in handling large-scale problems and the analytical solution it offers (Friston et al., 2007; Hoffman et al., 2013; Daunizeau et al., 2014; Zeidman et al., 2023). However, parameterized approaches such as VI are not without drawbacks, such as challenges in accurately approximating multi-modal posteriors and providing reliable uncertainty estimates.

In light of these limitations, efforts have been made to propose alternatives to variational inference for Bayesian model inversion in DCM (Chumbley et al., 2007; Aponte et al., 2016; Sengupta et al., 2015, 2016). A benchmark study was conducted using a comprehensive panel of non-parametric sampling-based methods on a neural mass model of event-related potentials (ERPs) measured in magneto/encephalography (MEG/EEG) recordings (Sengupta et al., 2015, 2016). These studies involved custom code for implementing the tested algorithms, which were released as Markov Chain Monte Carlo (MCMC) inference in the DMC reference toolbox Statistical Parametric Mapping (Friston et al. (1994); Penny et al. (2011)). Initial studies focused on gradient-free MCMC methods, including random walk Metropolis, slice-sampling, adaptive MCMC, and population-based MCMC with tempering (Sengupta et al., 2015). Gradient-free methods, although fast, struggled to thoroughly explore the posterior space and failed to produce a full, exploitable posterior distribution necessary for extracting key quantities, such as quantiles. Subsequent work focused on gradient-based MCMC methods, particularly emphasizing Langevin Monte Carlo (Roberts and Stramer, 2002; Girolami and Calderhead, 2011) and Hamiltonian Monte Carlo (HMC; Neal (2010)) algorithms (Sengupta et al., 2016). By leveraging the first-order gradients of the joint log-likelihood function, these classes of MCMC methods aim to alleviate the slow mixing problem and statistical inefficiency that plague gradient-free samplers. Despite tremendous effort and extensive testing, the results of previous studies (Sengupta et al., 2015, 2016) were not encouraging in terms of convergence and the number of independent samples produced per unit computational time. While fitting the data was relatively achievable within an acceptable error margin, obtaining accurate inference on the neurobiological parameters of the model proved to be more difficult. Gradient-based methods suffered from prohibitively high computational costs given the relatively small effective sample size they produced. Moreover, dynamics-based methods, such as HMC, face challenges in setting algorithmic parameters such as the step size and the number of steps for their numerical integration. The No-U-Turn Sampler (NUTS; Hoffman and Gelman (2014)) addresses these challenges by using a recursive algorithm to dynamically determine algorithmic parameters based on the geometry of the target distribution. Nevertheless, when dealing with nonlinear differential equations involving correlated parameters and partial observations, challenges related to achieving convergence within reasonable computational costs emerge.

This work aims to address the limitations encountered in previous approaches by exploring alternative programming languages and platforms for DCM implementation. It emphasizes the importance of cross-platform compatibility and accessibility in advancing neuroimaging research. Our research tackles the practical challenges of Bayesian inference in partially observable non-linear systems of differential equations, which are fundamental tools in most scientific domains. With advancements in computation, particularly in automatic differentiation and adaptive algorithms bolstered by the progress of machine learning techniques, the high computational costs of gradient-based methods are being significantly mitigated (Baydin et al., 2018). This development could potentially facilitate the routine application of sophisticated sampling schemes in DCM, and, broadly speaking, state-space ODEs models.

One promising solution lies in Probabilistic Programming Languages (PPLs), which are a class of programming languages designed to facilitate the specification, manipulation, and inference of complex probabilistic models (Gordon et al., 2014). These languages integrate standard programming constructs with probabilistic modeling, allowing for the seamless combination of deterministic and stochastic components. PPLs simplify the process of specifying complex probabilistic models. Instead of manually deriving mathematical formulations for inference, users can write models using high-level programming syntax, which makes the modeling process more intuitive and accessible. Additionally, PPLs provide built-in inference algorithms, saving time and reducing the potential for errors compared to manual implementation. Many PPLs are optimized for performance, leveraging advanced computational algorithms, techniques, and hardware acceleration to efficiently handle large datasets and complex models, ensuring scalability (Štrumbelj et al., 2024). PPLs such as Stan (Carpenter et al., 2017), PyMC (Abril-Pla et al., 2023), and Numpyro (Phan et al., 2019), provide powerful tools for probabilistic modeling, offering advantages in flexibility, automated inference, integration with computation, scalability, and applicability across diverse scientific domains.

We underscore the efficacy of PPLs, which, through implementation in various libraries with interfaces for common data science languages (such as Python and C++), achieve rapid fitting, high accuracy, and enhanced efficiency. Our results show significant improvements in the effective sample size obtained per unit of computational time, particularly by leveraging the JAX library (Bradbury et al., 2018), which offers composable function transformations and automatic differentiation for high-performance machine learning. Moreover, we conduct a comparative analysis between the NUTS and the variational methods. Through this comparison, we delineate the strengths and weaknesses of each method, ultimately showcasing their convergence to the same posterior distribution when successfully applied. We conduct comprehensive model comparison using both variational inference (assessed through free-energy; Friston et al. (2007); Penny (2012)) and MCMC frameworks (evaluated with fully Bayesian information criteria; Gelman et al. (2014a); Hashemi et al. (2021)). Our results demonstrate close alignment between methods in the MCMC framework and the free-energy obtained from variational inference, ensuring consistency in model comparison.

Importantly, another challenge for accurate DCM is the non-identifiability and its geometrical counterpart known as degeneracy, i.e., the existence of multiple equivalent solutions in the parameter space. Degeneracy is ubiquitous across biological systems (Tononi et al., 1999; Edelman and Gally, 2001), and crucial for the brain’s resilience and adaptability (Price and Friston, 2002; Sajid et al., 2020). While degeneracy provides an adaptive mechanism for maintaining brain function (degeneration and dedifferentiation; Fornito et al. (2015)), allowing compensatory strategies to deal with impairments and diseases, it also exacerbates the difficulties associated with non-identifiability in parameter estimation. This interplay contributes to multi-modality in the posterior distributions, where multiple sets of parameters can equally explain the observed data. This complicates the inference process and leads to wasted computational resources. We propose remedies for practical multi-modality, which manifests as convergence to different modes in repeated inference runs, overestimation or underestimation when using variational inference methods, or poor mixing in MCMC sampling. Our solutions include (1) optimal setting of algorithmic parameters, (2) initialization at the tail of prior distribution, and (3) weighted stacking of chains to average Bayesian predictive distributions (Yao et al., 2018), all of which significantly improve the inference.

In brief, our investigation underscores the balance between computational expense and inferential accuracy, highlighting the critical demand for flexible, efficient, and accurate Bayesian inference methods in neuroimaging. The tools are now accessible on the cloud platform EBRAINS (https://ebrains.eu), enabling users to explore more realistic brain dynamics underlying neurological conditions within a Bayesian causal framework.

## 2. Materials and methods

### 2.1. Dynamic Causal Modeling

Dynamic Causal Modelling (DCM; Friston et al. (2003); Razi et al. (2015); Frässle et al. (2017)) is a well-established statistical framework that allows for modelling causal interactions between brain regions, based on neuroimaging data, such as functional magnetic resonance imaging (fMRI), magnetoencephalography (MEG) or electroencephalography (EEG). It evaluates causal responses to diverse experimental manipulations, particularly on evoked responses to stimuli. Central to DCM is the concept of effective connectivity, which denotes the influence that one brain region or neuronal system exerts over another. It quantifies the directed interactions between distinct units, thereby delineating the causal relationships within the brain’s functional network. DCM formalizes the nonlinear dynamics of brain activity by representing them as a biologically informed nonlinear dynamical system (David et al., 2006b). Model inversion in DCM involves not only estimating the biophysical and effective connectivity but also enables source reconstruction through the inference of hidden states. In ERP models, these hidden states correspond to the activity in non-observed sources (David et al., 2006a).

### 2.2. Neural Mass Model

Neural Mass Models (NNMs; Deco et al. (2008); Coombes and Byrne (2018)) efficiently describe the collective behavior and interactions of large populations of neurons under the mean-field assumptions. They are widely used to understand brain dynamics at a macroscopic scale, during resting state (Spiegler et al., 2011; Rabuffo et al., 2021), task-related activities (Garrido et al., 2007; Chen et al., 2008), as well as in altered states such as anesthesia (Kuhlmann et al., 2016; Hashemi et al., 2018), healthy (Lavanga et al., 2023) and diseased conditions (Wang et al., 2023; Jirsa et al., 2023; Sorrentino et al., 2024).

In this study, we focused on a neurobiologically plausible generative NMM of event-related responses (ERPs) measured with MEG/EEG recordings (David et al., 2006b). The model comprises three interconnected neural populations (Figure 1**A**): spiny-stellate cells (*x*_1_), which receive the input current, inhibitory interneuron (*x*_7_), and the only observed population: excitatory pyramidal neurons (*x*_9_). The temporal evolution of model variables is described by a nine-dimensional system of ODEs (given by Eq. (1)), serving as a first-order approximation to delay differential equations by 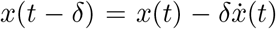. This evolution is governed by a set of ten parameters that are described in Table 1. The perturbation input to spiny-stellate cells is modeled as a Heaviside step function, where the parameter *u* represents its intensity (Sengupta et al., 2015, 2016):

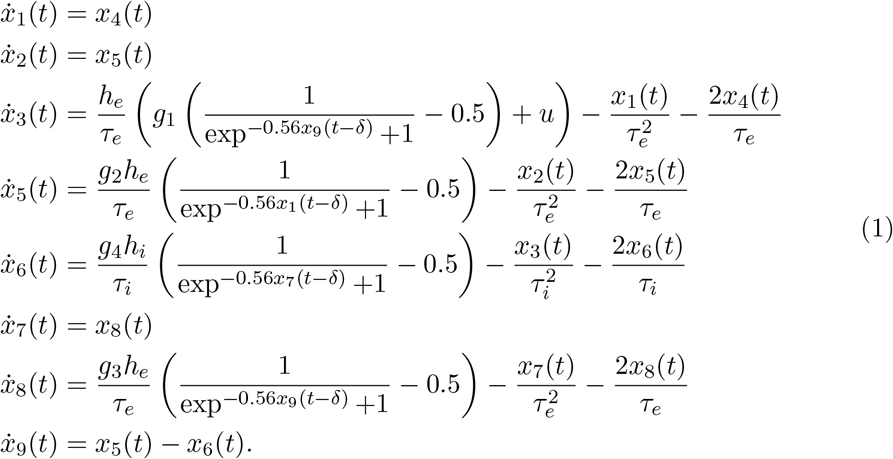

We simulate observations of pyramidal neuron voltage by (Euler-) integrating the model with a time step of *dt* = 0.1 msec (Figure 1**B**). A zero-centered Gaussian noise with a standard deviation of 0.1 is added to the observations, which are then downsampled by a factor of ten.

**Table 1.**
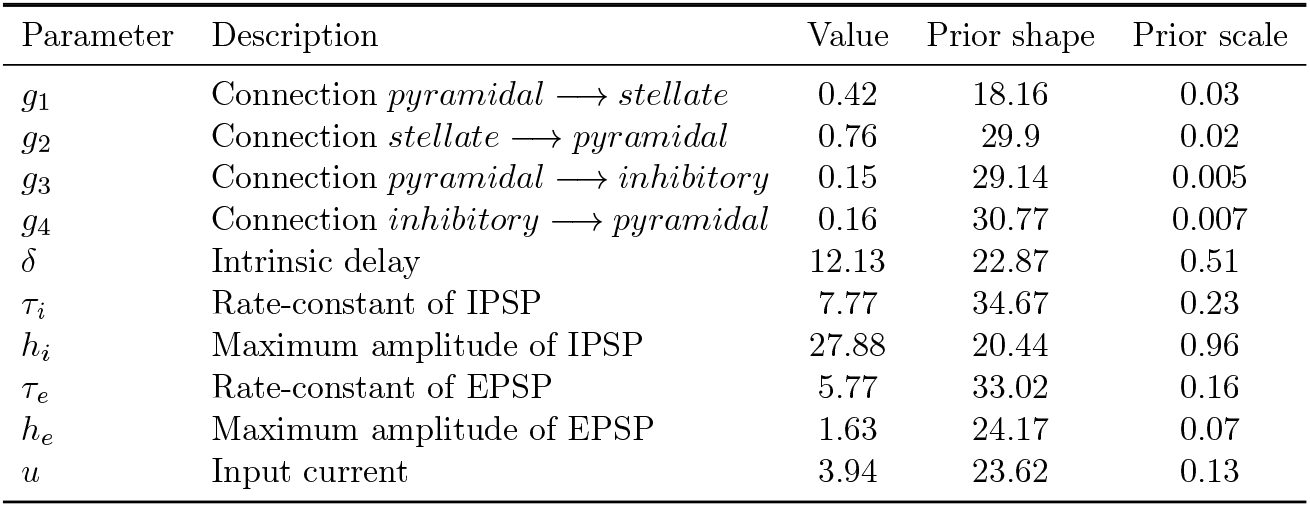
Parameters in a DCM of ERPs (David et al., 2006b) and their description. True values used to generate the data, along with the shape and scale of the Gamma prior placed for Bayesian analysis that ensure that about 50% of the sampled parameters result in unstable dynamics of the system (Sengupta et al., 2015, 2016). EPSP: Excitatory postsynaptic potentials; IPSP: inhibitory postsynaptic potentials.

| Parameter | Description | Value | Prior shape | Prior scale |
| --- | --- | --- | --- | --- |
| $g_1$ | Connection *pyramidal* → *stellate* | 0.42 | 18.16 | 0.03 |
| $g_2$ | Connection *stellate* → *pyramidal* | 0.76 | 29.9 | 0.02 |
| $g_3$ | Connection *pyramidal* → *inhibitory* | 0.15 | 29.14 | 0.005 |
| $g_4$ | Connection *inhibitory* → *pyramidal* | 0.16 | 30.77 | 0.007 |
| $\delta$ | Intrinsic delay | 12.13 | 22.87 | 0.51 |
| $\tau_i$ | Rate-constant of IPSP | 7.77 | 34.67 | 0.23 |
| $h_i$ | Maximum amplitude of IPSP | 27.88 | 20.44 | 0.96 |
| $\tau_e$ | Rate-constant of EPSP | 5.77 | 33.02 | 0.16 |
| $h_e$ | Maximum amplitude of EPSP | 1.63 | 24.17 | 0.07 |
| $u$ | Input current | 3.94 | 23.62 | 0.13 |

**Figure 1.**
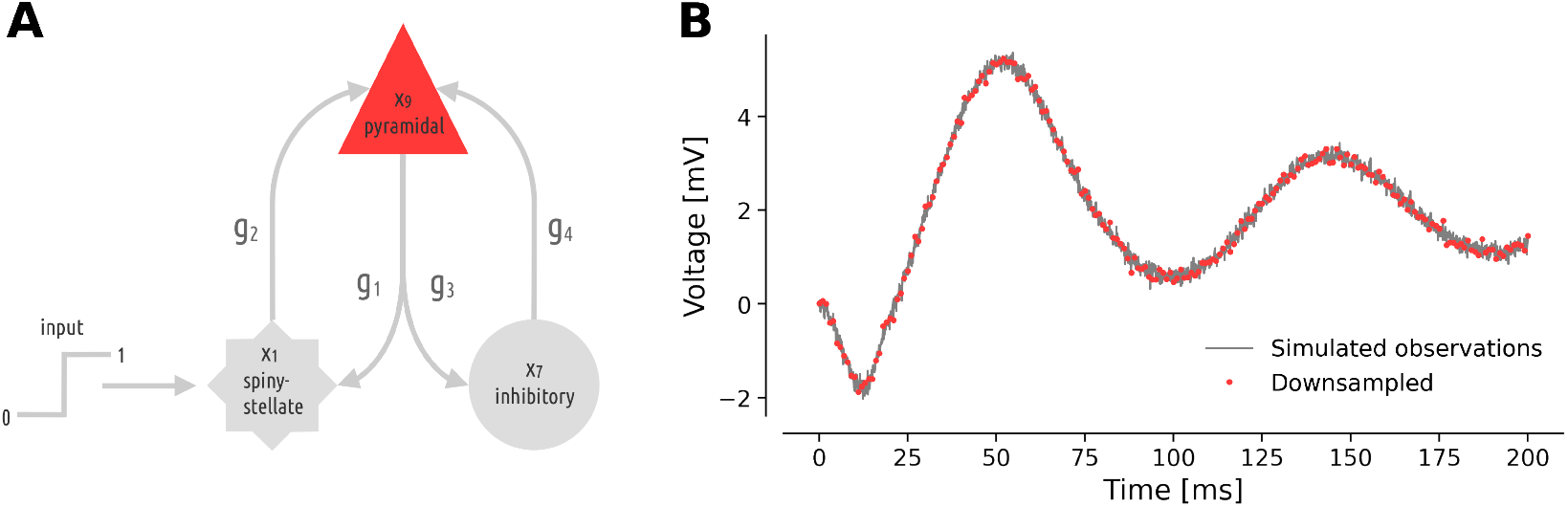
A DCM of ERPs (David et al., 2006b). (**A**) Schematic representation of the causal relations in the forward model consists of three neural populations, namely, pyramidal neurons, inhibitory interneurons, and spiny-stellate cells, driven by exogenous Heaviside input, according to Eq. (1). (**B**) The pyramidal cell voltage (in grey) is the only observable in the model, and was subjected to additive noise and downsampled by a factor of ten (in red).

### 2.3. Bayesian inference

Bayesian inference allows to take advantage of background knowledge about the parameters *θ* of a statistical model while updating this knowledge given new information provided by observed data *y*. The background knowledge, or prior, is encoded as a probability distribution *p*(*θ*), which reflects assumptions about the quantities to be inferred, before seeing the data. These assumptions are updated under the influence of the data, quantified by the likelihood *p*(*y* | *θ*), providing the posterior distribution *p*(*θ* | *y*), up to a normalizing constant:

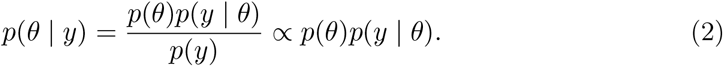

In this work, our goal is to infer the parameters of a DCM of ERPs given a simulated response of pyramidal neurons (see Eq. (1), and Figure 1). We assume a Gaussian likelihood and place Gamma priors parameterized according to Table 1 on model parameters.

### 2.4. Hamiltonian Monte Carlo

Markov Chain Monte Carlo (MCMC; Brooks et al. (2011)) methods are the gold-standard for generating samples from the posterior distribution by constructing a Markov chain that converges to this target distribution. Hamiltonian Monte Carlo (HMC; Duane et al. (1987); Neal (2010)) is the state-of-the-art MCMC algorithm that leverages deterministic Hamiltonian dynamics to inform Markov chains, enabling more efficient exploration of the target distribution. HMC introduces an auxiliary momentum parameter *ρ* to be jointly evolved with the position of model parameters *θ* in the parameter space, by integrating the Hamiltonian equations of motion given by:

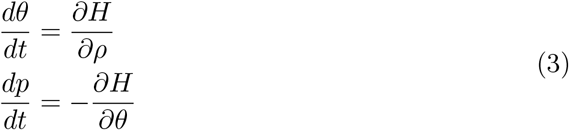

where *H*(*θ, ρ*) ≡−log *π*(*θ, ρ*) ≡−log *π*(*ρ θ*) − log *π*(*θ*) ≡ *K*(*ρ, θ*) + *V* (*θ*) is the Hamiltonian function. In this representation, *K*(*ρ, θ*) accounts for the kinetic energy and *V* (*θ*) for the potential energy. The leapfrog integrator intertwines half updates of momentum with full update of model parameters:

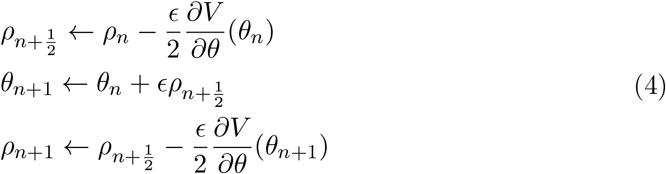

where *ϵ* is the step size. Outputs of the integration are then subjected to an accept-reject step (Gelman et al., 1995; Bishop, 2006). Hence, it is critical to monitor divergence, as it indicates that the numerical simulation of the Hamiltonian dynamics has failed, typically due to an energy increase suggesting a step size that is too large for complex geometries (Betancourt and Stein, 2011; Betancourt et al., 2017; Betancourt, 2017).

Although superior to gradient-free sampling algorithms, the performance of HMC is highly sensitive to the step size and the number of steps in the leapfrog integrator used to update the position and momentum variables in Hamiltonian dynamic simulation (Hoffman and Gelman, 2014; Betancourt, 2017; Betancourt et al., 2017). Its adaptive variant, the No-U-Turn Sampler (NUTS; Hoffman and Gelman (2014)), implements a recursive path to dynamically tune on-the-spot the hyper-parameters (step size and number of steps) of the symplectic integrator, thereby eliminating the need for hand tuning of these algorithmic parameters. For instance, the accept rate parameter (between zero and 1) is used to control the target acceptance probability for proposed steps during sampling, while the tree depth parameter controls the maximum number of leapfrog steps (specified as 2^*N*^).

The self-tuning sampler automatically adjusts trajectory length based on the behavior of the target distribution. However, due to numerical instabilities or highly curved and complex posterior geometries, the sampler may attempt to construct an infinitely long trajectory. To prevent this, a maximum trajectory length is imposed by setting the maximum tree depth parameter. Hamiltonian trajectories are numerically approximated by the leapfrog integrator, where the precision of this approximation is governed by the step size: for a fix length, a smaller step size results in more points along the numerical trajectory, thereby closely approximating the continuous trajectory, but at the expense of higher computational cost. Hence, the performance of HMC is contingent on the length of these trajectories. If the trajectory is too short, the sampler explores the parameter space inefficiently. On the other hand, if the trajectory is excessively long, the sampler may loop back to its starting point, leading to redundant exploration and the waste of computational time. Models with well-behaved geometry typically operate well with default hyperparameters values. However, for more complex models with intricate degeneracies, setting an appropriate proposal (or maximum) value for these hyperparameters necessitates careful consideration of the target distribution’s geometry to balance computational cost and accuracy (Hoffman and Gelman, 2014; Betancourt and Stein, 2011; Betancourt et al., 2017; Betancourt, 2017).

### 2.5 Variational Inference

Variational Inference (VI; Jordan et al. (1999); Wainwright et al. (2008)) is an alternative to MCMC methods that approximates probability distributions through an optimization problem, which typically results in much faster computation than MCMC methods. The core idea of VI is to posit a fixed-form (i.e., a parameterized) family of distributions and then to find the member of that family which is close to the target (Blei and Jordan, 2006; Blei et al., 2017; Zhang et al., 2018). In VI, closeness is expressed in the sense of the Kullback-Leibler (KL) divergence, which is an (asymmetric and non-negative) information-theoretic measure of proximity between two probability distributions. The goal is to minimize the KL divergence between the parameterized approximate posterior *q* (*θ*) ∈ 𝒬 from a family of tractable distributions and the true posterior *p*(*θ | y*). However, evaluating this divergence requires computing the evidence log *p*(*y*), which becomes intractable. Instead, VI optimizes an alternative objective, which (by the fact that *KL*(.) ≥ 0, or through Jensen’s inequality) forms a lower bound for log *p*(*y*), the marginal log likelihood of the observed data, known as the Evidence Lower Bound (ELBO; Neal and Hinton (1998); Paisley et al. (2012)) or negative free-energy (Friston, 2010; Zeidman et al., 2023):

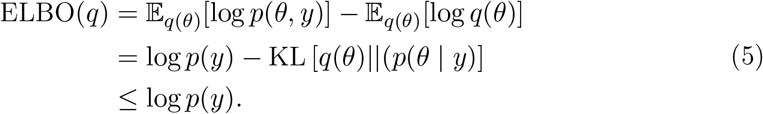

This allows us to perform approximate Bayesian inference by maximizing the ELBO with respect to *q*(*θ*), instead of directly minimizing the intractable KL divergence between the variational distribution and the true posterior.

In practice, VI transforms an inference problem into an optimization problem, requiring a full pass through the data at each iteration (Bishop, 2006). Stochastic Variational Inference (SVI; Hoffman et al. (2013)) defines local and global variational parameters and uses natural gradients in a stochastic optimization algorithm over subsamples (minibatches) of data to enhance the speed and scalability of variational approaches. In this study, we opted for SVI using Numpyro (Phan et al., 2019), a probabilistic programming library designed for fast, flexible, and scalable Bayesian modeling and inference. In addition to user-defined variational families, Numpyro supports several automatic guides that derive variational families from the generative model (Webb et al., 2019; Baudart and Mandel, 2021). Guides available in Numpyro include the mean-field and full-rank Gaussians, low-rank Gaussian and variational distributions parameterized by Normalizing Flows (such as Inverse Autoregressive Flow (Kingma et al., 2016) and Block Neural Autoregressive Flow (Cao et al., 2019)) that rely on the autoregressive property and transformations by neural networks to convert a simple distribution to any target.

In addition, Numpyro offers an automatic Laplace approximation. Laplace’s method approximates a function near its mode as a Gaussian density by leveraging its second-order Taylor expansion (quadratic approximation). The mean (mode) of the target Gaussian distribution is determined by the argmax of the function, while the covariance matrix is computed as the inverse of the negative Hessian matrix evaluated at the mode. Hence, the Laplace approach relies on the analytical solution, but to make it feasible, the log posterior needs to be twice-differentiable. Laplace variational inference, as the primary approach used in DCM (Friston et al., 2007; Daunizeau et al., 2014; Zeidman et al., 2023) leverages a quadratic approximation to the variational energy log *p*(*y, θ*) and defines a proposal quadratic approximated posterior *q*(*θ*) ≡ 𝒩 (*µ*, Σ^−1^), where

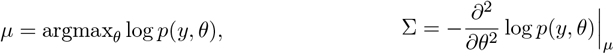

and gives the following expression for the ELBO under the Laplace approximation:

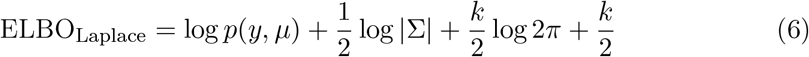

where *k* is the number of parameters and |.| denotes the determinant. In Numpyro’s implementation of variational Laplace in unconstrained space, a point estimate is inferred (maximum a posteriori, i.e., argmax_*θ*_ log *p*(*θ* | *y*), which is equivalent to argmax_*θ*_ log *p*(*y, θ*)) for the mean *µ*, by using Dirac delta distributions as (intermediate) variational family. The inverse of the Hessian is evaluated at this point, using JAX’s auto-differentiation for computing the Hessian. JAX (Bradbury et al., 2018; Frostig et al., 2018) is a Python library that provides a high-level tracer for implementing transformations (e.g. automatic differentiation, vectorization and JIT compilation) of Python functions. Numpyro wraps several JAX optimizers to be used in SVI. In this work, we use Adam (Kingma and Ba, 2014) with learning rate 0.0005.

Nevertheless, the model-specific derivation of VI requires a tedious and extensive effort to define a variational family appropriate for the probabilistic model, compute the corresponding objective function and gradients, and execute a gradient-based optimization algorithm. Automatic Differentiation Variational Inference (ADVI; Kucukelbir et al. (2016)) solves this problem automatically, in a black-box manner (Ranganath et al., 2014), without requiring analytic derivation or manual tuning by the user. ADVI is an automatic implementation of VI that derives an efficient and generic variational inference solution, requiring only a statistical model and dataset, without the need to specify a variational family or rely on Gaussianity or conjugacy assumptions (Kucukelbir et al., 2016). ADVI employs an unconstrained real-coordinate space mapping *θ* ⟼ *ζ* that facilitates the use of a single variational family for all models in a large class, implicitly generating non-Gaussian variational distributions in the original space. The mean-field implementation of ADVI posits a factorized Gaussian variational approximation,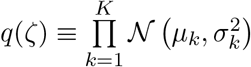, that is a product of Gaussians 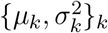, with variational parameters ignoring off-diagonal terms of the covariance (i.e., the degeneracy between parameters). The full-rank variant of ADVI defines *q*(*ζ*) ≡ 𝒩 (*µ, LL*^*T*^) with variational parameters *ϕ* = *{µ, L}* to be optimized, thereby capturing posterior correlations and producing more accurate estimates of marginal variances. In ADVI, the ELBO objective is approximated using Monte Carlo integration, and is to be maximized with stochastic (automatic differentiation-) gradient-based optimization (specifically, AdaGrad; Duchi et al. (2011)). In this work, we use the ADVI implementation (Kucukelbir et al., 2016) in the probabilistic software Stan (Carpenter et al., 2017).

### 2.6. Probabilistic programming languages

Probabilistic Programming Languages (PPLs; Gordon et al. (2014)) are a paradigm for automatic statistical modeling and machine learning that allows for the definition, manipulation, and inference of complex probabilistic models using high-level programming languages. They enable users to specify probabilistic models in a more intuitive and flexible manner, while leveraging high-performance computing and Automatic Differentiation (AD) through dependencies on specialized libraries. Central to the functionality of many PPLs is the concept of AD, a computational technique that efficiently and accurately calculates the derivatives of functions (Baydin et al., 2018). Unlike symbolic differentiation, which involves manipulating mathematical expressions, or numerical differentiation, which approximates derivatives using finite differences, AD breaks down functions into elementary operations and systematically applies the chain rule to compute derivatives. AD computes derivatives to machine precision, thereby avoiding the approximation errors inherent in numerical differentiation. It is notably efficient, with the computational complexity of obtaining gradients comparable to that of evaluating the function itself. AD plays a crucial role in enabling efficient and accurate inference using gradient-based methods in PPLs.

As of today, numerous Bayesian analysis software and packages offer embedded automatic differentiation and probabilistic sampling. While Stan has long been established in the statistical community, the rise of machine learning in the last decade has considerably boosted the development of automatic differentiation packages, paving the way for numerous gradient-based inference tools to be efficiently accessible. Although historically, statisticians have shown a penchant for the R programming language (for example, the R interface for Stan RStan was created in 2013, the Python interface PyStan in 2018), the grown popularity of Python within the scientific community has boosted the development of inference packages in this language. Pyro (Bingham et al., 2019) is a universal PPL based on PyTorch (Paszke et al., 2019). NumPyro (Phan et al., 2019) is a lightweight version of Pyro that runs on the JAX framework (Frostig et al., 2018; Bradbury et al., 2018), for efficient program optimization, parallelization, and GPU/TPU acceleration.

In this study, we implemented inference on a DCM of ERPs (see Eq. (1)) in several open-source PPLs from the top 20 most-downloaded packages related to Bayesian inference (see Štrumbelj et al. (2024), which also provides the list of R packages): CmdStanPy (Python interface to CmdStan (Lee et al., 2017)), PyMC (Abril-Pla et al., 2023), Numpyro (Phan et al., 2019) and BlackJAX (Cabezas et al., 2024). Although the choice of which PPL to use is at the convenience of the user, we benchmarked their efficiency, with a focus on the NUTS algorithm.

### 2.7. Model comparison and stacking

Model comparison involves evaluating and comparing different statistical models to determine which one best explains the observed data, taking into account both accuracy and complexity (Raftery, 1995). The criteria to assess and compare models balance model evidence, (or likelihood), that is, the fit to the data, and a penalty for model complexity, thus favoring models that achieve a trade-off between simplicity and explanatory power. Simplistic criteria like the Akaike Information Criterion (AIC; Akaike (1974), given by Eq. (7)) or its Bayesian variant (BIC; Schwarz (1978), given by Eq. (8)) are well-known and used according to a point estimation (Burnham and Anderson, 2002, 2004):

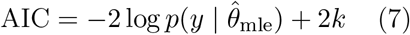

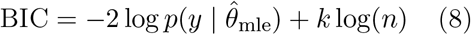

where log 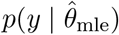 is the maximum log likelihood, *k* is the number of parameters in the statistical model, and *n* is the number of data points.

In the Bayesian context, where we have access to the full posterior distribution, other criteria have been proven to be more suitable (Gelman et al., 2014b; Hashemi et al., 2021). Based on expected log pointwise predictive density (elpd), the approximate Watanabe-Akaike Information Criterion (WAIC; Watanabe (2010, 2012); Vehtari et al. (2017), given by Eq. (9)) and cross-validated leave-one-out (CV-LOO; Vehtari et al. (2017); Gelman et al. (2014b), given by Eq. (12)), provide a more nuanced assessment of model fit and complexity.

The formula to compute the WAIC criterion is given by: (Gelman et al., 2014b)

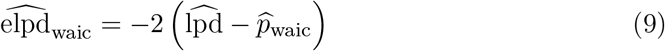

with the computed log pointwise predictive density 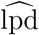 based on averaging with respect to posterior samples 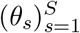, and simulation-estimated effective number of parameters 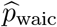, computed from the empirical variance given posterior samples,

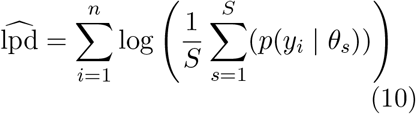

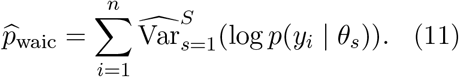

The Pareto-smoothed importance sampling approximation of CV-LOO (denoted by PSIS-LOO), which eliminates the need to perform actual leave-one-out cross-validation, is given by:

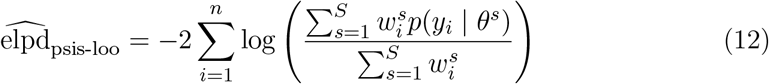

where 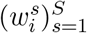 are Pareto-smoothed importance sampling weights (Vehtari et al., 2017).

In the DCM framework, the most popular inference algorithm, (Laplace) VI, is based on minimizing the ELBO (given by Eq. (5)), also known as the negative free-energy (Zeidman et al., 2023). This approach is widely used for model selection in DCM (Penny, 2012).

Related to model comparison, *stacking* to average Bayesian predictive distributions (Yao et al., 2018, 2020), weights and combines the distributions from multiple models (or chains) to provide more robust inference and enhance predictive performance. Given *K* models *M*_1_, … *M*_*K*_, the weights computed based on stacking of predictive distributions, are the solution of the optimization problem

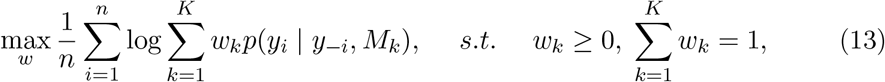

where *y*_−*i*_ is data without the *i*-th data point and the leave-one-out predictive density *p*(*y*_*i*_ |*y*_−*i*_, *M*_*k*_) can be estimated with 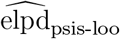 (Eq. (12)). Thanks to this approximation, readily implemented in the Arviz package (Kumar et al., 2019), the computation of stacking weights is fast, negligible compared to that of the inference part. We use stacking of predictive distributions to address post-hoc the discrepancies and multi-modality observed when running multiple chains or repetitions of inference on an ill-defined model. It is worth noting that the stacking approach is not limited to outputs of MCMC algorithms, and can be applied as well to VI (Yao et al., 2018, 2020).

### 2.8. Convergence diagnostics

After running an MCMC sampling algorithm, it is necessary to conduct statistical analysis to evaluate the convergence of the chains. The Gelman-Rubin statistic 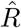 (Gelman and Rubin, 1992), which approaches 1 under optimal convergence, is the standard estimator to verify the within- and between-mixing of multiple Markov chains. Effective sample size (ESS; Vehtari et al. (2021)) is another useful indicator that quantifies the uncertainty relative to the estimation of posterior quantities such as quantiles from the typically auto-correlated draws from Monte-Carlo Markov chains (Geyer, 2011). A strong sampling efficiency in the core of the posterior is characterized by a high bulk ESS, which is computed for rank normalized values using split chains (Vehtari et al., 2021). The sampling efficiency in the tails of the posterior is defined as the minimum of the ESS for the 5% and 95% quantiles.

More generally, in Bayesian analysis, useful quantities for detecting issues in inference include posterior z-scores, computed for each model parameters, which are a combination of bias and precision of the posterior, 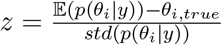, and should be close to 0 in the ideal case (Betancourt, 2017; Hashemi et al., 2020).

All experiments were conducted on a Linux machine with a 3.60 GHz Intel Core i7-7700 CPU (4 cores) and 32 GB of memory.

## 3. Results

In this section, we present our findings on various strategies and comparisons aimed at improving and evaluating Bayesian inference methods for a DCM of ERPs (see Eq. (1) and Figure 1). We begin by outlining three strategies for addressing multi-modality in inference: fine-tuning the hyperparameters of the sampler, leveraging initialization, and employing stacking techniques based on predictive accuracy. Following this, we provide a comparison between VI family and NUTS in fitting the ODE model, highlighting the strengths and limitations of each approach. We then conduct Bayesian model comparison under both variational and MCMC frameworks, ensuring consistency in model evaluation. Lastly, we benchmark the sampling efficiency of three popular PPLs (namely, Numpyro, PyMC and Stan), showcasing the impact of programming environment on inference performance.

### 3.1. Multi-modality

Multi-modality is often encountered in biological system (Edelman and Gally, 2001), when distinct sets of parameter values can explain the data equally well. In MCMC, running multiple Markov chains with diverse starting points is a standard practice to ensure the reliability of inference. This approach helps in assessing convergence and ensures that the chains have explored different regions of the parameter space. By comparing the results of multiple chains, one can determine if they are converging to the same region or if there are significant discrepancies indicating multiple modes. It can also happen that multi-modality is not present in the true data generating process, but rather is a result of the inference process. When the fit to the data or the posterior predictive checks are inadequate, it indicates issues such as a misspecified model, data quality concerns, or problems with the inference method. Regarding the latter, ensuring that the algorithm used to estimate the posterior does not get stuck near a local optimum is crucial.

We encounter discrepancies in the results of repeated inference (e.g., multiple HMC chains) on the DCM of ERPs (see Figure 1 for the generated observations). We perform a first round of inference on the NMM model (Eq. (9)) for which we set diffuse Gamma priors (see Supplementary, Table S1), illustrating the multi-modality that arises during the inference process (Figure 2**A**). The chains, ran with random initial conditions, can fail to recover the system dynamics (Figure 2**B**), and are therefore not indicative of true multi-modality in the data generating process, rather they are misleading the inference. Nevertheless there could exist other modes capable of generating accurate dynamics in observation (see Figure S1). The chains that show non-stationary patterns are discarded, however several chains show convergence to a stationary distribution that nevertheless does not reflect the true data generating distribution (Figure 2**C**). These results indicate the challenges in achieving convergence to the true data generating distribution, especially given a high-dimensional ODE model and diffuse priors.

**Figure 2.**
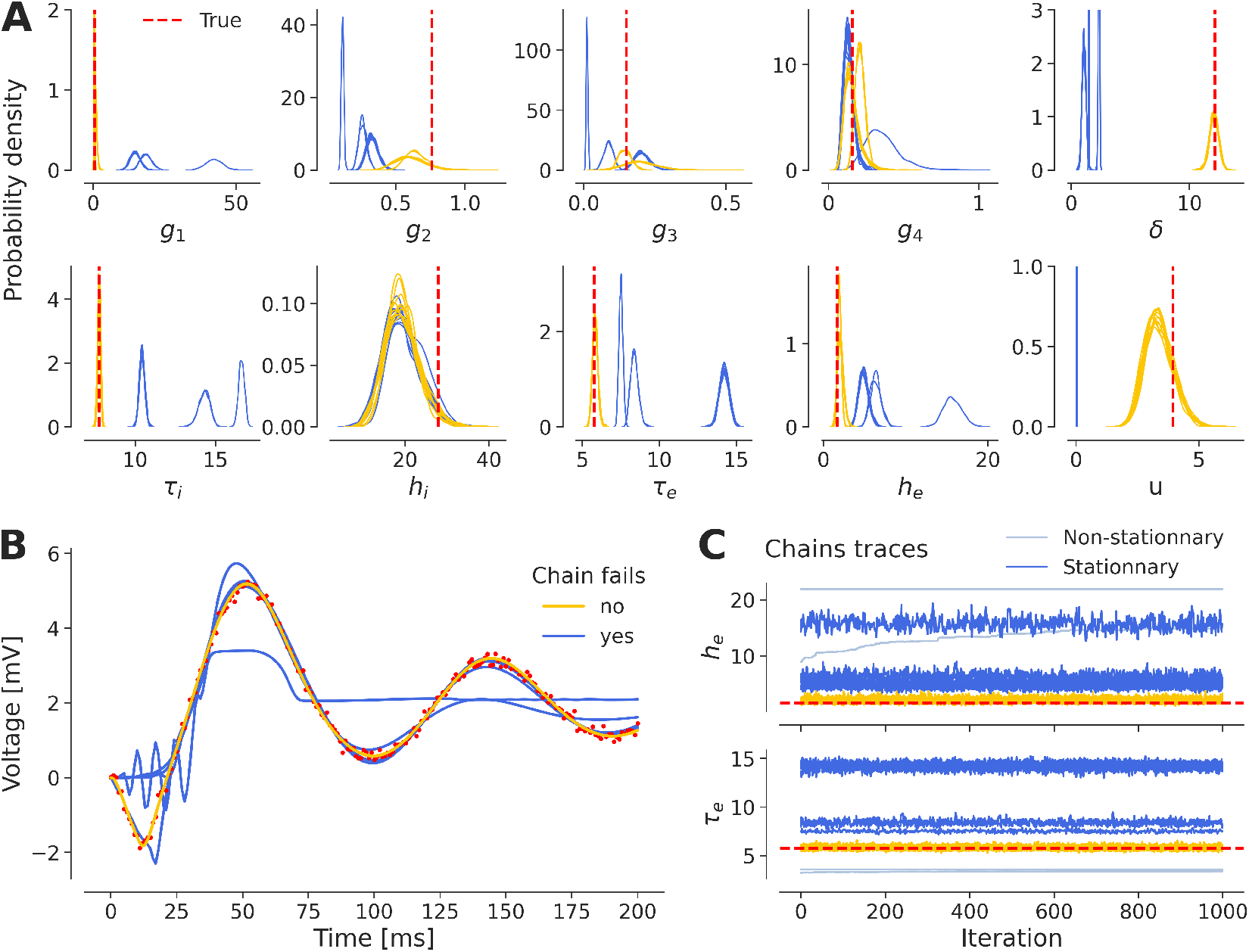
Multi-modality arise in the inference process. (**A**) The estimated posterior distributions depending on their closeness (in blue and yellow), to true values (displayed in red) targeting different regions or modes of the parameter space. (**B**) Chains that fall into local optima (in blue) in the parameter space fail also to recover perfectly the true dynamics of the system (in red), contrary to the chains that target the correct posterior mode (in yellow). (**C**) Chains that show non-stationary patterns are discarded (in light blue), however a chain can converge to a stationary distribution that is a local solution (in blue) rather than the global optimum (in yellow).

Placing an informative prior has been proposed as a solution to mitigate the issues of multi-modality and degeneracy (Hashemi et al., 2021; Baldy et al., 2023). In the rest of this paper, we use a set of weakly informative Gamma priors motivated by Sengupta et al. (2015, 2016) as described in Table 1. We note that enforcing even more informative priors does not guarantee to solve the issues of multi-modality, as we have shown in Supplementary, Figure S2, and we need to find other solutions to address the exploration issues encountered by the inference scheme. It is important to note that this lack of convergence is not a flaw specific to MCMC sampling (here, NUTS) and using other methods such as variational inference result in the same issue (see Supplementary, Figure S3).

As the first step, we aim to improve the convergence of the algorithm by enhancing the performance of the integrator (see Figure 3). Adjusting the hyper-parameters of the inference algorithm, such as the step size and the number of leapfrog steps in HMC, may improve exploration in the search space. Adaptive versions of these algorithms, such as NUTS, can automatically tune these parameters during the warm-up phase. However in NUTS, other important parameters include the maximum tree depth of the binary tree representing the evolution of the integrator, ensuring finite-time integration if the No-U-Turn termination criterion is never met. Additionally, the target acceptance rate of Hamiltonian trajectories submitted in a Metropolis step influences the integration step size.

**Figure 3.**
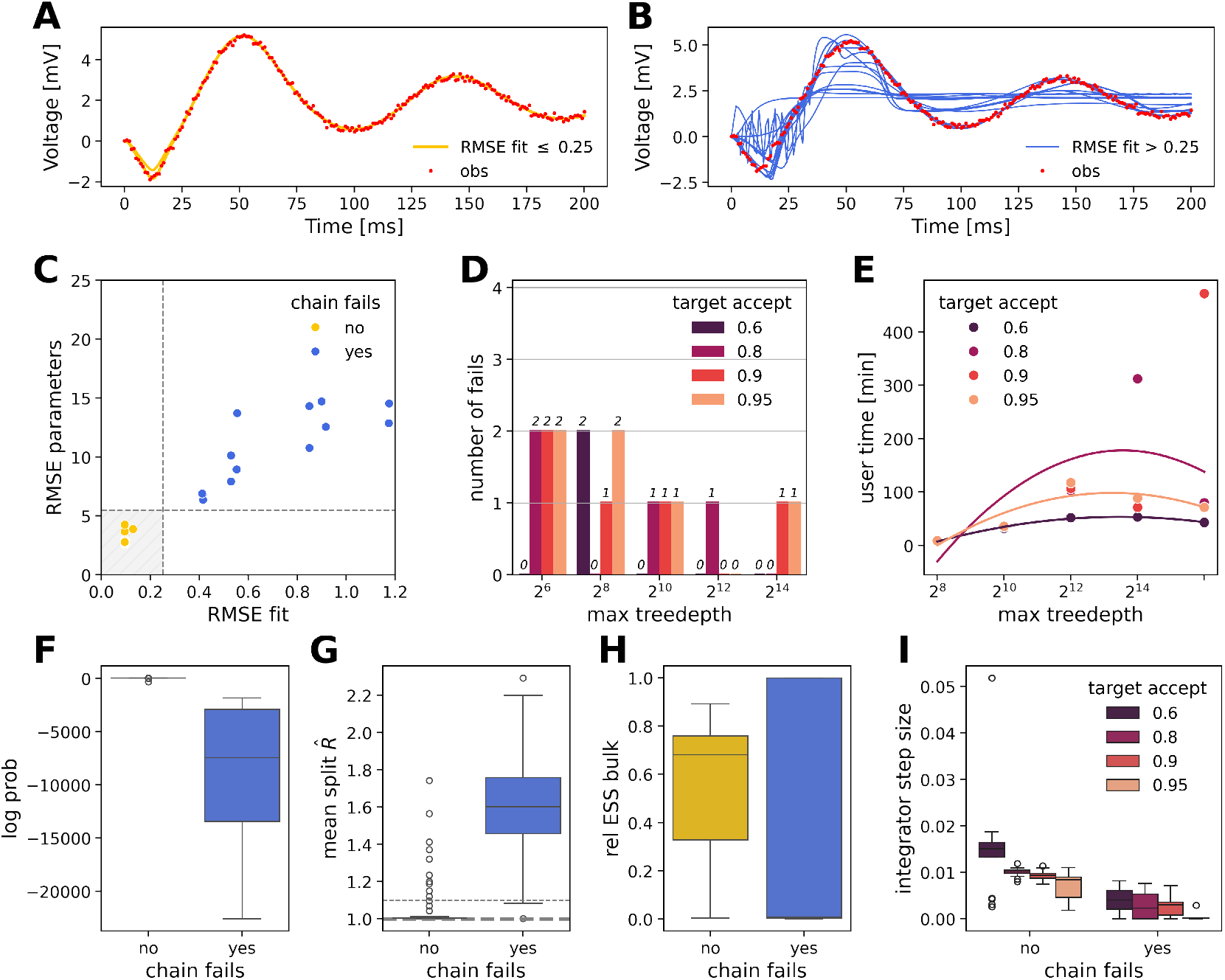
Fine-tuning hyper-parameters of the No-U-Turn Sampler. (**A**) Observations (in red) and faithful fits (*RMSE* ≤ 0.25, in yellow). (**B**) Observations (in red) and inaccurate fits (*RMSE >* 0.25, in blue). (**C**) Estimation error in retrieving the model parameters (RMSE parameters) versus the estimation error in retrieving the observation (RMSE fit). There is a clear relationship between these two quantities and successful inferences (in yellow) are located within the shaded grey area. (**D**) Count of chains that fail to recover the data, out of 4 parallel chains, with respect to 20 different NUTS hyperparameter settings. (**E**) Computational time of NUTS (with 1000 warmup and 1000 samples) with respect to hyperparameter settings. (**F**) Log probability, (**G**) Mean split 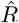, computed with the two halves of a chain as a measure of within-chain convergence and (**H**) Relative effective sample size (ESS) in the bulk of the posterior. (**I**) Step-size of the leapfrog integrator with respect to target acceptance rate.

Here we demonstrate the results of fine-tuning the hyperparameters of NUTS to ensure both within-chain and between-chain convergence. In Figure 3, we summarize the inference results obtained for 20 combinations of hyper-parameters, in terms of fit to the data (RMSE fit), closeness of estimated parameters from the ground truth (RMSE parameters), log probability (as a measure of prediction for the observed data), and convergence indicators, such as relative ESS, and split 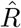 (see section 2).

Figure 3**A**,**B** illustrates the observations and successful overlapping fits (Figure 3**A**, with *RMSE* ≤ 0.25), or failed recoveries of the observation (Figure 3**B**, with *RMSE >* 0.25), out of a total of 80 chains (20 different hyperparameters configurations, run with 4 chains each). From Figure 3**C**, we observe that a poor fit to the data is closely related to a poor estimation of model parameters. In Figure 3**D**, we reported the number of chains that fail to recover the data and ground truth parameters, for each tested pair of hyperparameters (target acceptance and max tree depth). For this specific model and data, a tree depth of 2^8^ nodes is sufficient to prevent premature termination of Hamiltonian integration but is not a guarantee of sufficient exploration that achieves convergence. Setting the target acceptance rate to 0.6 solved the convergence issues (Figure 3**D**).

In the face of challenging convergence, the computation time does not follow a strict law and heavily depends on a chain being stuck in a local minimum (Figure 3**E**). Chains that are able to faithfully fit the data show a significantly higher log probability (Figure 3**F**), as well as better within-chain mixing and convergence (see split 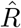; Figure 3**G**). However, there is no clear difference in effective sample size (Figure 3**H**), except that chains that fail in the task of recovering the data or ground truth parameters can achieve either extremely low or extremely high ESS. Setting a low target acceptance rate enforced exploration and enabled larger integration steps (Figure 3**I**). It also shows that in this model, successful fits are characterized by enhanced exploration, with, on average, larger step sizes. These results indicate the critical effect of algorithmic parameters on efficient and accurate inference, while their hand-tuning for convergence is painstaking work and does not easily generalize.

As the second step, we aim to improve convergence by leveraging the information provided by the weakly informative priors (see Table 1). An alternative to the tedious tuning of algorithmic parameters is initializing the chains in a non-zero probability region of the prior. Initializing chains with a random sample taken uniformly from the tails of the prior (specifically below the 2.5th and above the 97.5th percentile, constituting a 5% probability region) ensured 100% convergence, regardless of the values of HMC hyperparameters (but within reasonable ranges), as shown in Figure 4. For the observation (Figure 4**A**), and initialization strategy at the tails (Figure 4**B**), we can see that in this case, all the chains converged successfully without any failures, out of a total of 80 chains (Figure 4**C, D**). In such a regime where convergence is straightforward, the computation time scales quadratically with the maximum tree depth (Figure 4**E**). The log probability (Figure 4**F**), the split 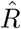 (Figure 4**G**) and the relative ESS (Figure 4**H**) confirm the convergence of HMC chains. The influence of the target acceptance rate on the adaptation of the step-size of the leapfrog integrator is again highlighted in Figure 4**I**. Large integrator steps are achieved (Figure 4**I**), in agreement with the corresponding panel in the previous figure (Figure 3**I**, where the chains that did not fail demonstrated a larger step-size on average compared to the ones that failed), favoring exploration through proposals over exact integration. These results demonstrate that initializing the chains at the tail of the prior provides a very straightforward and efficient convergence for our DCM model.

**Figure 4.**
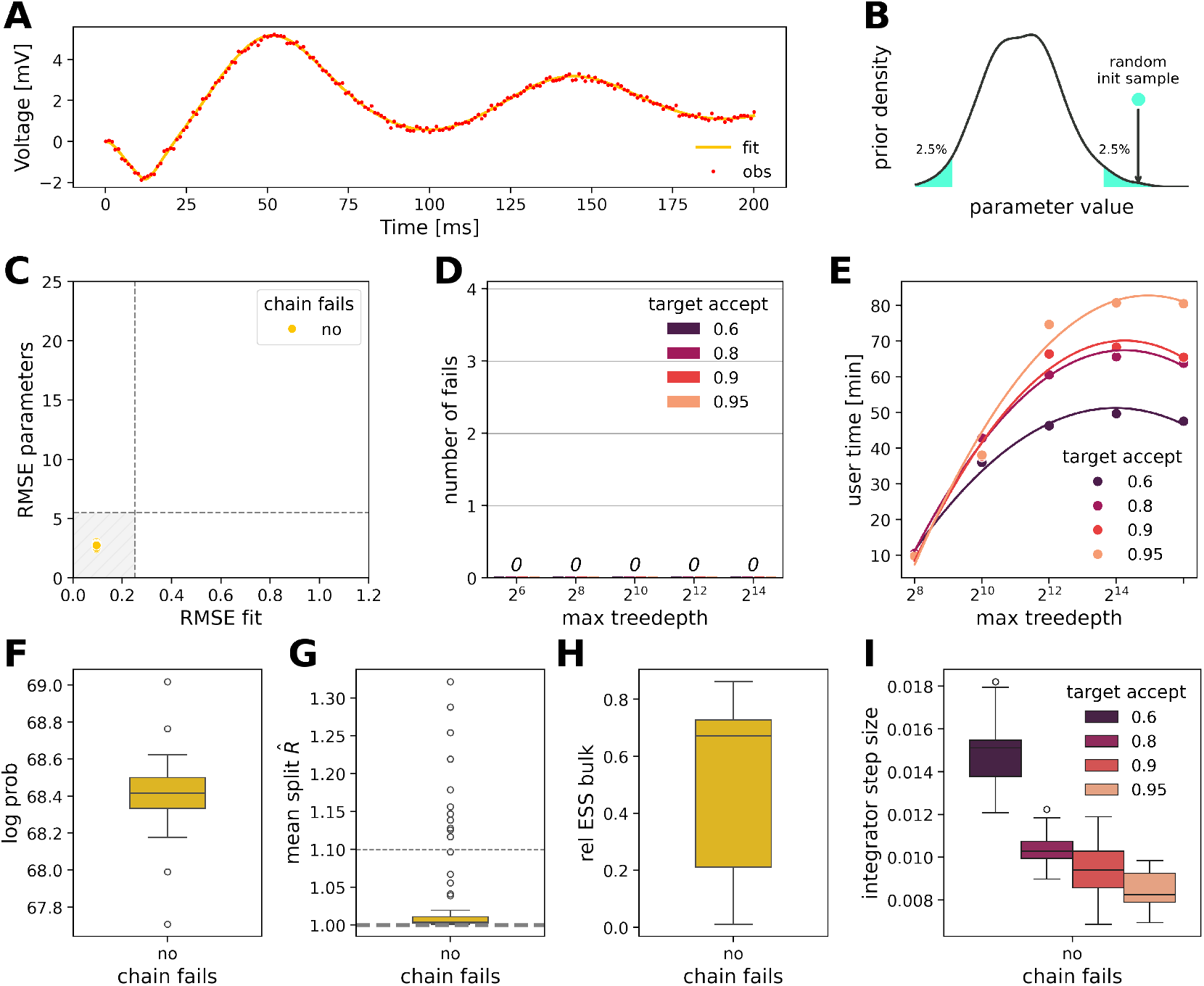
Initialization of the chains in the tails of the prior as a remedy for convergence issues. (**A**) All of the 80 chains successfully recover the observed data. (**B**) Initialization scheme: the initial position of a chain in parameter space is set to a random sample taken uniformly in the tails of the priors. (**C**) Estimation error in retrieving the model parameters (RMSE parameters) versus the estimation error in retrieving the observation (RMSE fit). All initialized runs are successful inferences and are located within the shaded grey area. (**D**) No initialized chain, out of a total 80 chains, fails to recover the data, run with 20 different NUTS hyperparameter settings. (**E**) Computational time of NUTS (with 1000 warmup and 1000 samples) with respect to hyperparameter settings. (**F**) Log probability, (**G**) Mean split 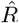, computed with the two halves of a chain as a measure of within-chain convergence and (**H**) Relative effective sample size (ESS) in the bulk of the posterior. (**I**) Step-size of the leapfrog integrator with respect to target acceptance rate.

As the third step, we aim to propose a more generic solution for achieving chain convergence without manual tuning of algorithmic parameters or relying on the priors. This solution involves running standard practice randomly initialized chains, and subsequently stacking them post-hoc (Yao et al., 2018, 2020). This is achieved through weighted averaging based on their respective predictive power (see Eq. (13)).

In Figure 5, we illustrate an example of mismatched inference where chains end up in different regions of the parameter space, some exhibiting Dirac-delta type distributions (i.e., strongly centered around a single point in the parameter space, indicating very low variability in the estimates of parameters). We can see that, unlike the naive pooling approach, stacking the chains successfully smooths out local artifacts and phantom modes (Figure 5**A**). Stacking to average Bayesian predictive distributions resulted in a perfect reproduction of the dynamics in the data, unlike pooling, which still suffers from errors induced by failed chains (Figure 5**B, C**, respectively). It is worth noting that the stacking approach can also be applied to outputs of variational inference with multiple initializations (Yao et al., 2018, 2020).

**Figure 5.**
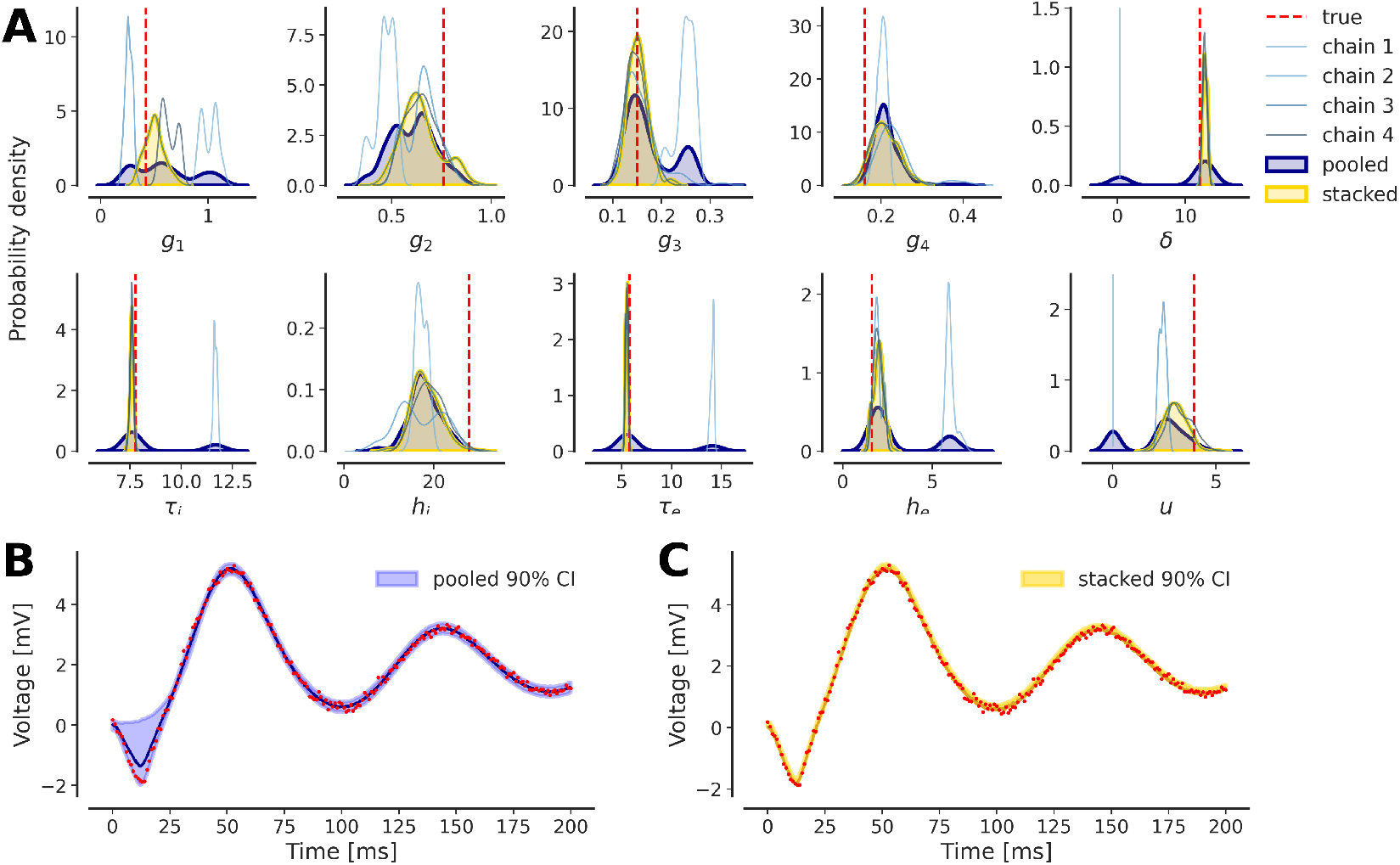
Addressing multi-modality in inference with stacking. (**A**) Markov chains (in shades of blue) have converged to different distributions that arise from different modes in the parameter space. (**B**) Pooling the chains (in dark blue) does not filter out artifact modes that do not perfectly reproduce system dynamics and data. (**C**) Stacking (in yellow) applies weights based on the predictive distribution of each chain and successfully smooths out local artifacts, resulting in better predictive performance.

In sum, these results indicate that achieving reliable and robust Bayesian inference for ODE models requires understanding the intricacies of the model and its parameters, and careful monitoring of convergence, even when relying on automated and state-of-the-art algorithms. Although at first glance, the model seemed to produce inconsistent results, we were able to stabilize inference using three methods: the tedious one– hand tuning the algorithm, the simplest yet very effective one– initialization in the prior, and the more general approach– stacking the chains.

### 3.2. Algorithmic benchmark

Here we compare the performance of HMC versus VI methods. More specifically, we compare NUTS and (mean-field and full-rank variants of) ADVI, as well as the Laplace approximation, with initialization at the tails of the prior to ensure convergence. For consistency, we ran four parallel chains with NUTS (200 warmup and 200 sampling iteration, maximum tree depth of 10 and acceptance rate of 0.8), and four repetitions of variational inference (with 10^5^ iterations and a single ELBO sample).

In terms of computational time, the four NUTS chains using Stan took around 52 minutes, compared to 4 minutes using NumPyro. The four repetitions of mean-field and full-rank ADVI using Stan took around 19.4 min and 18 min, respectively, in total (see Table 2 for exact values). Using Numpyro, they took 3.22 min and 12.98 min, respectively. These results demonstrate 2-13 times faster computational time using NumPyro for NUTS or ADVI. The automatic Laplace approximation proposed in Numpyro was the fastest of all methods, taking only 3.12 min total for the four repetitions of 10^5^ iterations. Hence, using NUTS or Laplace in NumPyro, we can sample the posterior in 3-4 min.

**Table 2.** Benchmarking of self-tuning variant of HMC, the No-U-Turn Sampler (NUTS), versus variational inference methods in solving an ODE model given by Eq. (1), using Stan and Numpyro implementations.

| Software | Algorithm | Time [sec] | Mean $\hat{R}$ | RMSE fit | Min. corr. | Max. corr. | Min. z-score | Max. z-score |
| --- | --- | --- | --- | --- | --- | --- | --- | --- |
| Stan | Mean-field | 1164.259 | 1.221 | 0.104 | 0.002 | 0.738 | 0.957 | 69.561 |
|  | Full-rank | 1080.777 | 1.006 | 0.104 | 0.015 | 0.985 | 0.115 | 1.260 |
|  | HMC-NUTS | 3115.238 | 1.007 | 0.104 | 0.003 | 0.981 | 0.006 | 2.374 |
| Numpyro | Mean-field | 193.583 | 1.062 | 0.009 | 0.004 | 0.234 | 1.067 | 266.635 |
|  | Full-rank | 779.127 | 1.003 | 0.097 | 0.002 | 0.922 | 0.422 | 19.579 |
|  | Laplace | 187.758 | 1.002 | 0.097 | 0.004 | 0.982 | 0.150 | 2.475 |
|  | HMC-NUTS | 245.1952 | 1.009 | 0.097 | 0.002 | 0.982 | 0.243 | 2.560 |

Figure 6 displays the posterior samples obtained by each algorithm for a few selected parameters (see Supplementary, Figure S4 for pairplot of all parameters).

**Figure 6.**
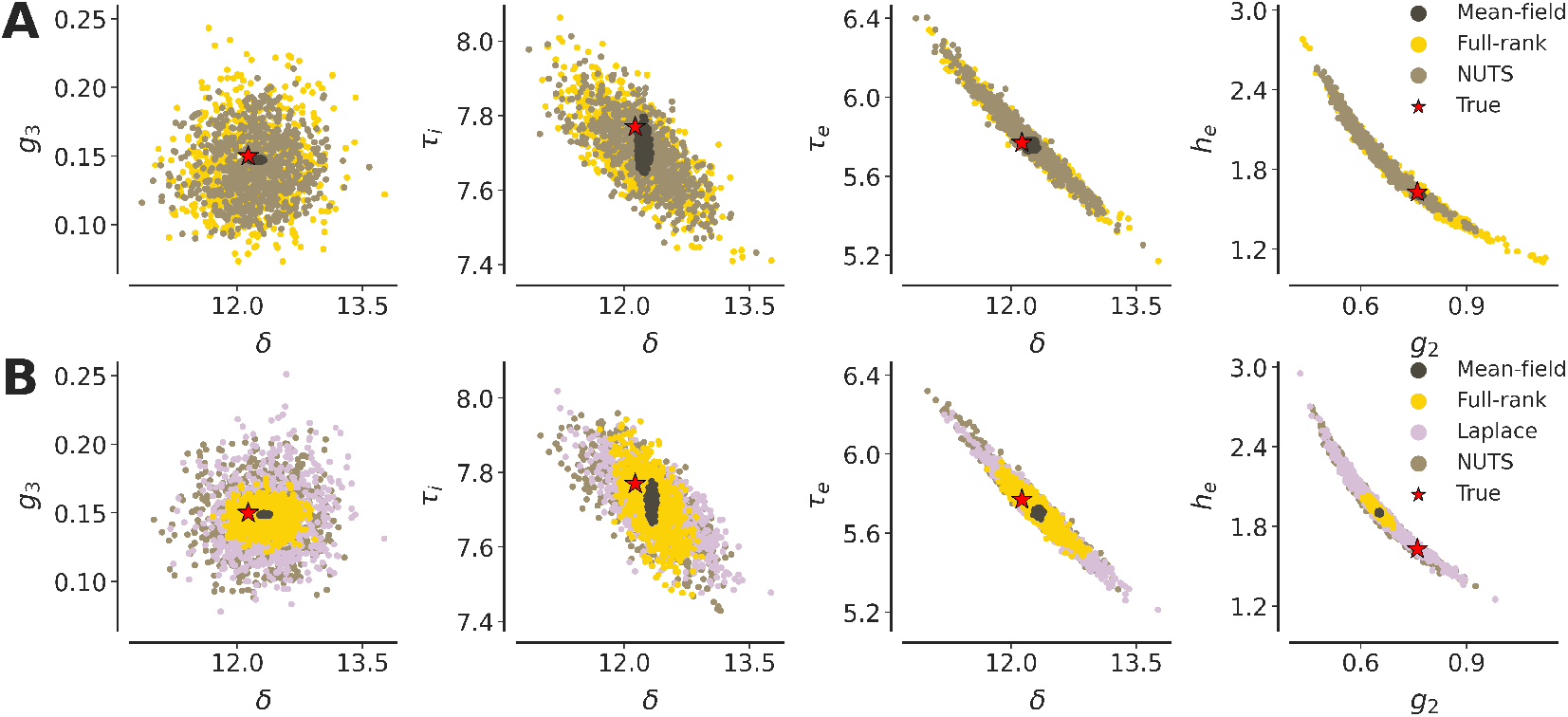
Pairplots of posterior samples obtained with variational inference methods (mean-field ADVI: dark brown, full-rank ADVI: yellow, automatic Laplace approximation: mauve) and HMC sampling (NUTS: light brown), for a selected subset of model parameters, using (**A**) Stan and (**B**) Numpyro. NUTS and the Laplace approximation both accurately provide posterior samples that encompass the true generative parameters (in red) and capture complex geometries in the posterior. In both softwares, the mean-field method underestimates marginal variances and produces uncorrelated and under-dispersed outputs. Numpyro’s full-rank ADVI provides overconfident posteriors which can overlook the true parameter value.

Indicators of convergence and accuracy, the mean 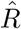 accross all parameters (although only interpretable for HMC), RMSE of the fit, minimum and maximum pair-wise correlations, and z-scores are reported along with computational time in Table 2. The NUTS, (Stan’s) Full-rank ADVI, and (Numpyro’s) Laplace approximation, all show satisfactory convergence.

In terms of accuracy, the results indicated that, as expected, (Stan’s) full-rank ADVI performs equivalently to NUTS in capturing complex posterior geometries (e.g., see the pairplot for *g*_2_, and *h*_*e*_ in Figure 6) and the true generative parameters, although it produces slightly more diffuse posterior distributions. Numpyro’s implementation of the full-rank Gaussian variational family gave significantly less dispersed posteriors than its Stan counterpart, at times over-confidently overlooking ground truth parameters (see the paiplot for *g*_2_, and *h*_*e*_ in Figure 6). The Laplace approximation offered by NumPyro yields estimated posterior distributions that are equivalent to those obtained using NUTS. A known limitation of mean-field ADVI is the systemic underestimation of marginal variances (Kucukelbir et al., 2016), which we also observe for the model under study, in both Stan and NumPyro implementations. The mean-field algorithm provides uncorrelated and under-dispersed outputs, which fail to capture several of the ground truth model parameters.

In sum, these results indicate the accuracy and efficiency of estimation using NUTS and the Laplace approximation (the later implemented only in NumPyro), with both algorithms producing reliable posterior distributions in 3-4 minutes.

### 3.3. Model comparison

This section focuses on model comparison, as a central tool within DCM. We generated observations using a full model of the system (see Figure 1 and Table 1). Subsequently, we fit an alternative model with removing connectivity between certain regions (see Figure 7).

**Figure 7.**
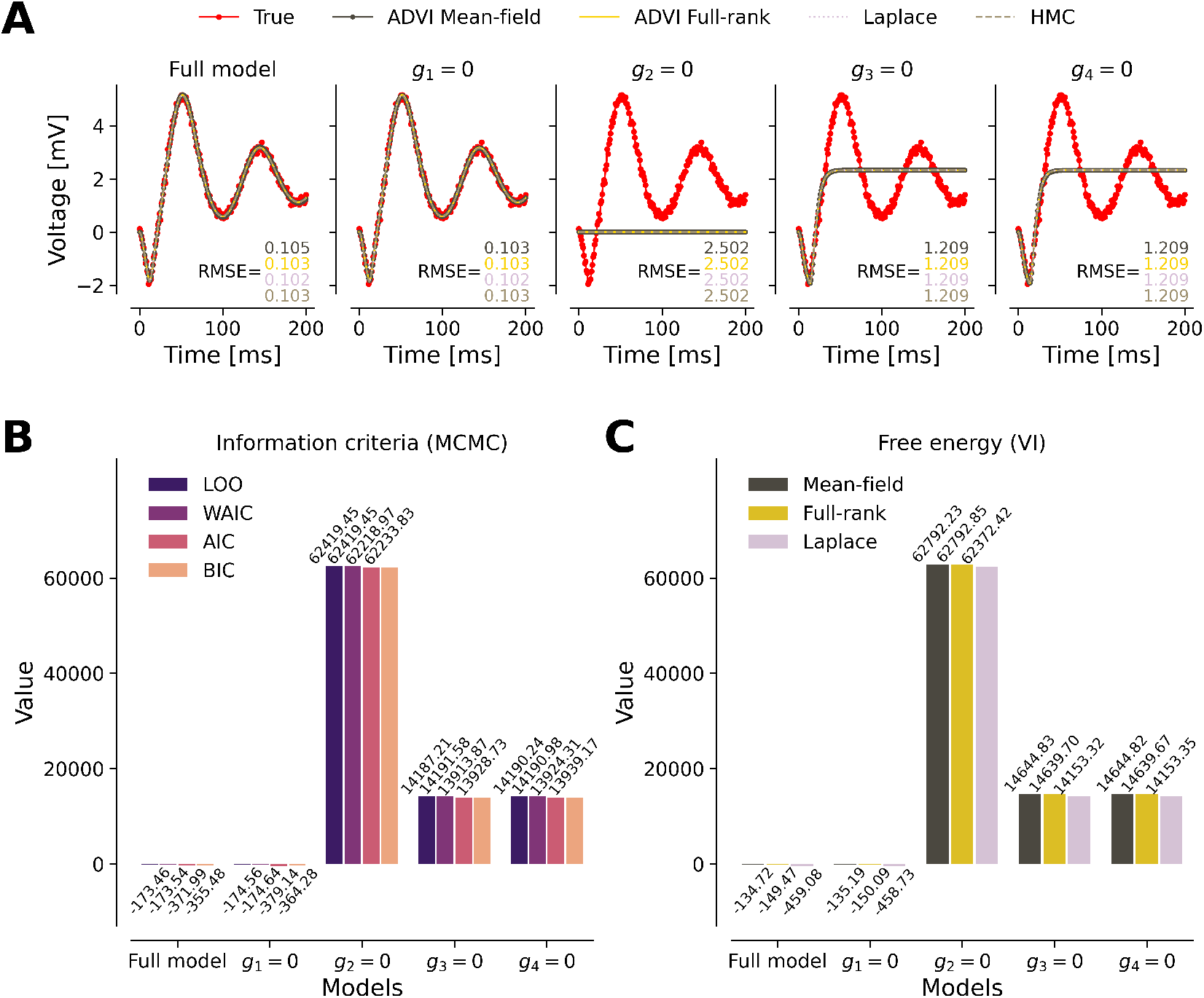
A complete taxonomy of model comparison methods. The full-model is compared to subset models that each set one connectivity parameter to zero. (**A**) Only the full model and the model where *g*_1_ = 0 are able to recover the dynamics of the system. (**B**) Model comparison criteria (AIC, BIC, WAIC, LOO) for each model fitted with MCMC (here the NUTS algorithm). (**C**) Free-energy as a model comparison criterion in variational inference (here the mean-field and full-rank ADVI, and automatic Laplace algorithms). All criteria indicate that the reduced model with *g*_1_ = 0 is superior, while the model with *g*_2_ = 0 performs the worst. The smaller the value of criteria, the better the model’s predictive ability.

We observe that the full model and the model setting *g*_1_ = 0 –thus vanishing the strength of connection from pyramidal neurons to spiny-stellate cells– both perfectly fit the observation. However, setting one of *g*_2_ = 0 (connection from spiny-stellate to pyramidal cells), *g*_3_ = 0 (connection from pyramidal to inhibitory cells) or *g*_4_ = 0 (connection from inhibitory to pyramidal cells) results in large deviation from the observation. More precisely, setting *g*_2_ = 0 leads to a worse fit, whereas *g*_3_ = 0 and *g*_4_ = 0 result in similarly poor fits (see Figure 7**A**, and the RMSE values).

We note that parameters for which estimation is most affected by the cuts in connections are the intrinsic delay *δ*, and rate constants of the membrane *τ*_*i*_ and *τ*_*e*_. Their posterior distribution estimated by ADVI and NUTS, with respect to each model, are shown with that of all other parameters in Supplementary, Figure S6.

Model comparisons based on classical AIC (Eq. (7)), and BIC (Eq. (8)) as well as Bayesian WAIC (Eq. (9)), and LOO cros-validation (Eq. (12)) in the MCMC frame-work (Figure 7**B**) align closely with free-energy obtained from variational inference (Figure 7**C**). Our model comparison results indicate that the model with *g*_1_ = 0 is the best, while the model with *g*_2_ = 0 performs the worst (Figure 7**B, C**). Interestingly, setting parameter *g*_1_ = 0 causes no loss in fitting the observation performance compared to the full model. Hence, the model comparison favorites such reduced model, as it diminishes by one the number of fitted parameters. Note that the smaller value of comparison criteria indicates the better model’s predictive ability.

Although this is straightforward in the definition of AIC/BIC, as the penalty term depends explicitly on the number of parameters, WAIC and LOO incorporate a more intricate calculation. They take into account the effective number of parameters (variance of the log-likelihood, given by Eq. (11)), which depends on the structure and complexity of the model and the influence of the prior distributions. This leads to a more accurate reflection of both the model fit and its generalization capabilities, ensuring that the selected model is not only well-fitted to the current data but also robust for predicting new, unseen data. Notably, the fully Bayesian information criterion WAIC closely approximate LOO cross-validation.

### 3.4. PPLs benchmark

This section focuses on fitting the model using NUTS within several popular Python-based PPLs. We compare their efficiency in terms of computational cost and quality of the inference obtained (convergence of chains and effective sample size). To ensure a fair comparison across PPLs, we ran NUTS with identical settings in each: default NUTS parameters (max tree depth of 10, acceptance rate of 0.8), four parallel chains, initialization in the tails of the prior, and 200 warmup and 200 sampling iterations.

Our implementation reveals that NumPyro and PyMC (using the BlackJAX back-end) were approximately 16 times faster than Stan (Figure 8**A** and Table 3) achieving virtually identical convergence properties, 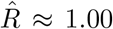 and effective sample sizes (see Figure 8**B, C** and Table 3). The only notable difference was Numpyro achieving a lower effective sample size in the bulk of the posterior (ESS bulk), that is in the high probability region, compared to the two other samplers (Figure 8**B** and Table 3).

**Table 3.** Benchmarking of NUTS implementation in solving an ODE model given by Eq. (1) in several PPLs. NUTS was run with default parameters in each software (maximum tree depth 10 and target acceptance rate 0.8) with 200 warmup and 200 collected samples.

| Package | Time [sec] | Mean ESS | | Rel. ESS | | Mean ESS bulk / sec | Mean $\hat{R}$ | Divergences |
| --- | --- | --- | --- | --- | --- | --- | --- | --- |
|  |  | bulk | tail | bulk | tail |  |  |  |
| NumPyro | 55.532 | 508.890 | 491.538 | 0.636 | 0.614 | 9.164 | 1.004 | 0 |
| PyMC x BlackJAX | 59.076 | 592.603 | 512.673 | 0.741 | 0.641 | 10.031 | 1.005 | 0 |
| CmdStanPy | 993.862 | 596.246 | 492.626 | 0.745 | 0.616 | 0.600 | 1.006 | 0 |

**Figure 8.**
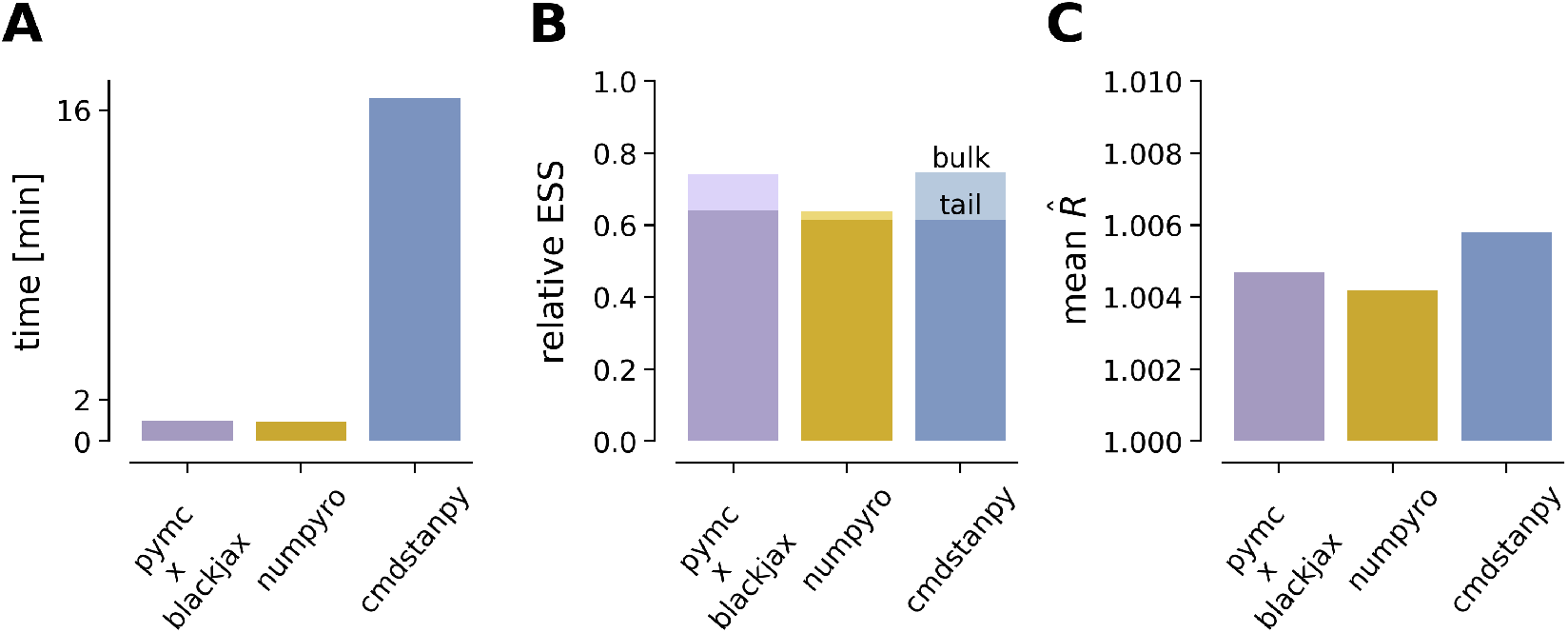
Benchmarking of NUTS implementation in solving an ODE model given by Eq. (1) across different popular PPLs. We report the following metrics for our comparison: (**A**) Computational time in minutes (four parallel chains, 200 warmup steps, and 200 samples). (**B**) Relative effective sample size ESS, i.e., number of effective samples divided by total draws. (**C**) Mean value of 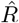 across all model parameters (measure of convergence between and within chains in MCMC, closer to 1 indicates better convergence).

## 4. Discussion

### 4.1. Related work

In evaluating the performance of various MCMC methods for Bayesian model inversion in DCM, previous research highlighted significant challenges (Sengupta et al., 2015, 2016). The gradient-free methods investigated in Sengupta et al. (2015) included random walk Metropolis, slice-sampling, adaptive MCMC, and population-based MCMC with tempering. Among these, slice sampling emerged as the most efficient gradient-free algorithm in terms of effective samples, yet it still provided less than 10 effective samples out of 1400 draws (after excluding 600 burn-in samples from an initial 2000). Gradient-based methods, including Hamiltonian Monte Carlo (HMC) and Langevin Monte Carlo (LMC) with two variants–Langevin diffusion on a Euclidean manifold (LMC-E) and a Riemannian manifold (LMC-R)–also faced substantial difficulties (Sengupta et al., 2016). Specifically, LMC-E yielded an effective sample size of only 11 out of 14000 samples (after excluding 6000 burn-in samples from an initial 20000), while HMC produced fewer than 100 effective samples despite an extensive run time of 45 hours. The computing performance of HMC in this case was voluntarily not optimized to assert for worst-case behaviour.

By leveraging advanced computational techniques and optimized automatic differentiation libraries, HMC can achieve efficient and accurate inference, overcoming the obstacles faced by earlier studies. This provides a more encouraging perspective on the use of Monte Carlo methods in DCM. In this work, we achieved MCMC convergence of 4 parallel chains using only 200 burn-in samples, and obtained accurate inference in less than 1 minute (see Table 3). The implementations in Numpyro, PyMC, and Stan enabled us to perform very efficient inference with NUTS, achieving an effective sample size that was more than half of the actual number of samples. We note that the slower performance of Stan compared to the other PPLs lies in the non-vectorized Euler integration loop, which accounts for most of the computational time. A more efficient implementation using vectorization could potentially improve the inference in Stan compared to others. While we observed that Numpyro or PyMC (using BlackJAX or Numpyro backends, as shown in Supplementary Figure S7, and Table S2) had a computational advantage over Stan with this specific model, it is important to note that this advantage should not be generalized. Our objective was not to conduct a comprehensive benchmark of PPLs across different inference tasks. Previously, the computational efficiency of Stan and Numpyro has been compared across several datasets (Baudart et al., 2021), with results showing that while Numpyro tends to outperform Stan across numerous datasets (around 2.3x speedup), there are instances where Stan performs better. Thus, the performance difference can vary depending on the specific dataset and task.

The final point to mention here is that PyMC provides implementations of a large number of gradient-free and gradient-based MCMC algorithms, as well as Sequential Monte Carlo (SMC; Doucet et al. (2001); Sisson et al. (2007)) algorithms. Consistent with previous studies (Sengupta et al., 2015), our results also indicated that NUTS, as gradient-based MCMC algorithm, significantly outperformed gradient-free algorithms such as the Metropolis family (see Supplementary Figure S7). SMC outperformed the aforementioned gradient-free samplers and was significantly faster than NUTS. However, its performance heavily depends on the acceptance threshold value.

### 4.2. Degeneracy

Degeneracy, the ability of structurally different elements to perform the same function or yield the same output, is a ubiquitous property in biological systems, and evident at genetic, cellular, system, and population levels (Edelman and Gally, 2001). Degeneracy is a fundamental aspect of neural architecture and is crucial for the brain’s resilience and adaptability (Sajid et al., 2020). By allowing multiple neural circuits or brain regions to compensate for each other, degeneracy ensures that the brain can maintain functionality despite damage or perturbations (Fornito et al., 2015; Wang et al., 2024). This (many-to-one mapping) capability is essential for processes such as learning, memory, and recovery from injury, where different neural pathways can adapt and take over functions when primary pathways are compromised. However, while degeneracy provides a robust mechanism for maintaining brain function, it poses challenges for inference, such as deciphering a bijective (one-to-one mapping) relationship. The overlapping functionalities of different neural elements can lead to difficulties in pinpointing specific causal pathways and accurately estimating model parameters. This complexity underscores the need for advanced computational methods and robust probabilistic approaches to disentangle the contributions of various neural components and better understand the intricate dynamics of brain function.

The overlapping functionalities inherent in degeneracy add another layer of complexity to the issue of non-identifiability in modeling neural dynamics. Structural non-identifiability (Raue et al., 2009; Wieland et al., 2021; Hashemi et al., 2023) is a common issue in partially observed dynamical systems described by ODEs. For a model to be structurally identifiable, it must allow for a unique set of parameters for any given output, ensuring that the solution to the inverse problem is not only unique but also depends continuously on the data. This well-posedness is crucial from a statistical viewpoint, as it enables robust parameter estimation, even with minor data perturbations (Latz, 2020). Given the often imprecise nature of data, we should expect these estimates to remain relatively stable with small changes. This stability is crucial for reliable inference and model validation. However, ill-posedness arises when the reconstruction of causes from observed effects becomes unstable, meaning that vastly different causes could produce similar effects. This poses significant challenges for accurately identifying causal relationships in complex systems. Structural identifiability issues stem from the model’s inherent design, which can render some parameters indeterminable (see Figure S8, where, in the present case, the connectivity parameter *g*_1_ shows clear structural non-identifiability). In contrast, practical identifiability challenges occur when the available measurements lack sufficient information to determine the parameters with the required precision. In the present model, profile likelihood methods (Wieland et al., 2021; Raue et al., 2011, 2009) suggest the practical non-identifiability of input parameter *u* beyond a certain threshold (see Figure S8). In inference, degeneracy manifests as a lack of information, complicating the extraction of precise and reliable parameter estimates (as we observed in Figure 2, where inference converges to several modes of parameter space, failing to recover the underlying dynamics of the data). The stacking of predictive distributions from multiple MCMC chains that we employed as a remedy (see Figure 5) leverages their combined information, thereby enhancing the overall information content and addressing the challenges posed by degeneracy. Initializing chains within prior ranges (see Figure 4) is another solution that incorporates additional information, effectively mitigating the degeneracy encountered during inference on ODE models.

### 4.3. Variational Inference

The mean-field approximation, while computationally efficient, is known to underestimate marginal variances (Kucukelbir et al., 2016), as we also observed in our results for a DCM of ERPs (see Figure 6). The simplification inherent in the meanfield approach can lead to overly confident parameter estimates. In contrast, full-rank approximation mitigates this issue by capturing correlations between variables, thus providing a more accurate representation of the posterior distribution. Similarly, the Laplace approximation, when applied without the mean-field assumption, has shown impressive results, producing posterior inferences that are competitive with those obtained via HMC. Based on analytical derivations, this method combines advantageous computational efficiency with robust inferential accuracy, making it a viable alternative to more computationally intensive techniques like HMC. Note that, in our comparisons, we ran the Laplace VI with 10^5^ iterations to ensure consistency in the benchmark with other algorithms. However, the Laplace method converges quickly, and its computational time can be significantly reduced through early stopping, without compromising the quality of the results.

The potential downsides of the Laplace approximation are due to its purely local nature, depending solely on the curvature of the posterior around the optimum (Zhang et al., 2018). Still, the most common advice when working with difficult posterior geometry is to reparametrize the model to obtain a log convex posterior (Betancourt et al., 2017; Jha et al., 2022). The Laplacian approximation may still therefore be widely applicable. A second computational constraint of the Laplace approximation is the need to invert the Hessian matrix of second-order derivatives, which has complexity *O*(*n*^3^) with the number of parameters. As with posterior geometry, reparametrization can allow structuring the Hessian into mean and full rank components, to sidestep costly large matrix inversions.

One of the core advantages of ADVI is that it removes the need for manual tuning by the user. In ADVI, the main idea is to represent the gradient as an expectation, and to use Monte Carlo techniques to estimate this expectation. Although ADVI leverages Gaussian mean-field and full-rank distributions, it operates with transformed variables, which induce non-Gaussian variational distributions in the original variable space. Ultimately, ADVI aims at enabling a generic inference algorithm, functioning as a black-box, without requiring analytical computation of the ELBO or assumptions about the variational family.

Beyond these methods, traditional variational inference typically minimizes the variational lower bound using the KL divergence (Penny et al., 2003; Kiebel et al., 2008). However, recent advancements have explored the use of alternative divergence measures, such as the Rényi divergence (Van Erven and Harremos, 2014; Li and Turner, 2016; Sajid et al., 2022). This approach introduces a hyperparameter that, when set to one, reduces to the KL divergence. By adjusting this parameter, the approximation can shift between mode-seeking behavior, which focuses on identifying the highest probability regions (but includes variability and is not reduced to point estimates), and mass-covering behavior, which aims to cover the entire distribution more comprehensively. We note that the ELBO objective leveraging Rényi divergence is implemented in Numpyro. Investigations into these types of alternatives in variational inference methods or inference networks (Rezende and Mohamed, 2015; Kingma et al., 2016, 2019) were beyond the scope of this study and remain to be explored in future research.

### 4.4. Model comparison

We conducted a comprehensive model comparison in both VI and MCMC frameworks. Variational approximation typically relies on maximizing the ELBO, which equivalently minimizes the free-energy, to assess model performance and compare different models. The ELBO (or negative free-energy) provides an estimate of model evidence that balances model fit and complexity. In the MCMC framework, model comparison works in an analogous way and is based on quantities computed post-hoc and without additional cost (Vehtari et al., 2017; Gelman et al., 2014b), such as classical AIC (Akaike, 1974) and BIC (Schwarz, 1978), or the more recently developed WAIC (Watanabe, 2010, 2012) and LOO (Vehtari et al., 2017), which are tailored to the Bayesian context (Gelman et al., 2014b). Our results showed that model comparisons based on AIC, BIC, WAIC, and LOO in the MCMC framework aligned closely with free-energy from variational inference (Figure 7). For researchers accustomed to variational inference, this finding provides reassurance that model comparison can be seamlessly transitioned to the MCMC framework without losing consistency in evaluation.

By definition, classical information criteria (AIC/BIC) are agnostic to prior information as they rely on maximum likelihood estimation, and the penalty term is determined solely by the number of parameters and observed data. In contrast, fully Bayesian information criteria (WAIC) and LOO cross-validation enable the incorporation of prior information into the model’s fit to data, thereby improving out-of-sample prediction accuracy through the integration of (clinical or biological) knowledge. This approach has been used in medical applications such as identifying epileptogenic zones in the brain (Hashemi et al., 2021) or predicting drug plasma concentration in alcohol use disorder (Baldy et al., 2023). Notably, the Bayesian LOO expected log point density approximates classical LOO cross-validation, which avoids the need to repeat the fitting process, making it a powerful and efficient estimate especially in contexts where fitting models can be computationally expensive. Moreover, WAIC closely approximates Bayesian LOO, indicating a direct link between information theory and cross-validation (Vehtari et al., 2017; Gelman et al., 2014b).

### 4.5. Limitations

This work focused on synthetic data to validate the algorithms rather than directly validating the model against empirical data. We limited our investigation to a single neural mass model, which may not fully capture the complexities of more elaborate DCM models involving intricate neural networks at larger scales. Extending our analysis to whole-brain level DCM presents a significantly more complex inversion problem that may necessitate the incorporation of sparse priors or other regularization techniques to obtain meaningful results. Our current work focused on ODE state-space models, but estimating dynamic and observation noise for fMRI data poses additional challenges for convergence. Investigating MCMC and variational inference methods for the DCM family operating at higher-dimensional model configurations subject to noise remains an area of future research.

### Information Sharing Statement

All code is available on GitHub (https://github.com/ins-amu/DCM_ERP_PPLs) and in Ebrains collaboratory drive (https://drive.ebrains.eu/).

## Acknowledgements

This research has received funding from EU’s Horizon 2020 Framework Programme for Research and Innovation under the Specific Grant Agreements No. 101147319 (EBRAINS 2.0 Project), No. 101137289 (Virtual Brain Twin Project), and government grant managed by the Agence Nationale de la Recherche reference ANR-22-PESN-0012 (France 2030 program). We thank Matthieu Gilson, Andrea Brovelli, and Daniele Marinazzo for fruitful discussions. The funders had no role in study design, data collection and analysis, decision to publish, or preparation of the manuscript.

## Author contributions

Conceptualization: V.J., and M.H. Methodology: N.B., M.W., and M.H. Software: N.B, M.W., and M.H. Investigation: N.B., and M.H. Visualization: N.B., Supervision: V.J. and M.H., Funding acquisition: V.J. Writing - original draft: N.B., and M.H. Writing - review & editing: N.B, M.W., V.J, and M.H.

## 5. Supplementary

**Table S1.** Parametrization of the diffuse and informative Gamma priors placed on model parameters in preliminary analysis.

| Parameter | Diffuse prior |  | Informative prior |  |
| --- | --- | --- | --- | --- |
|  | shape | scale | shape | scale |
| $g_1$ | 5 | 0.25 | 60 | 0.007 |
| $g_2$ | 5 | 0.25 | 250 | 0.003 |
| $g_3$ | 5 | 0.1 | 150 | 0.001 |
| $g_4$ | 2 | 0.1 | 80 | 0.002 |
| $\delta$ | 24 | 0.5 | 120 | 0.1 |
| $\tau_i$ | 34.67 | 0.23 | 110 | 0.07 |
| $h_i$ | 20.44 | 0.96 | 180 | 0.15 |
| $\tau_e$ | 33.02 | 0.16 | 110 | 0.05 |
| $h_e$ | 25 | 0.1 | 85 | 0.02 |
| $u$ | 23.62 | 0.13 | 130 | 0.03 |

**Table S2.** Algorithmic benchmarking in PyMC.

| Metric | Time [sec] | Mean ESS |  | RMSE params | RMSE fit |
| --- | --- | --- | --- | --- | --- |
|  |  | bulk | tail |  |  |
| Slice Sampler | 7.318314 | 46.583626 | 60.231617 | 9.838069 | 71.760229 |
| Metropolis | 4.649500 | 12.202948 | 28.362368 | 0.796275 | 5.467409 |
| DE MetropolisZ | 5.095385 | 12.190079 | 6.533076 | 0.967134 | 2.051315 |
| DE Metropolis | 6.521602 | 10.428617 | 30.754647 | 0.868549 | 0.102877 |
| SMC e=0.1 | 6.521602 | 10.428617 | 30.754647 | 0.868549 | 0.102877 |
| SMC e=10 | 798.342054 | 699.147368 | 4.140204 | 1.029628 | 0.531720 |
| NUTS Pytensor | 20.620693 | 22.501308 | 19793.222673 | 2.684353 | 0.169526 |
| NUTS Nutpie | 5.732669 | 68.935616 | 2052.947144 | 1.420048 | 5.771916 |
| NUTS Blackjax | 555.987747 | 480.066809 | 84.007686 | 0.850000 | 0.131545 |
| NUTS Numpyro | 623.706941 | 527.442519 | 84.043669 | 0.850000 | 0.131903 |

**Figure S1.**
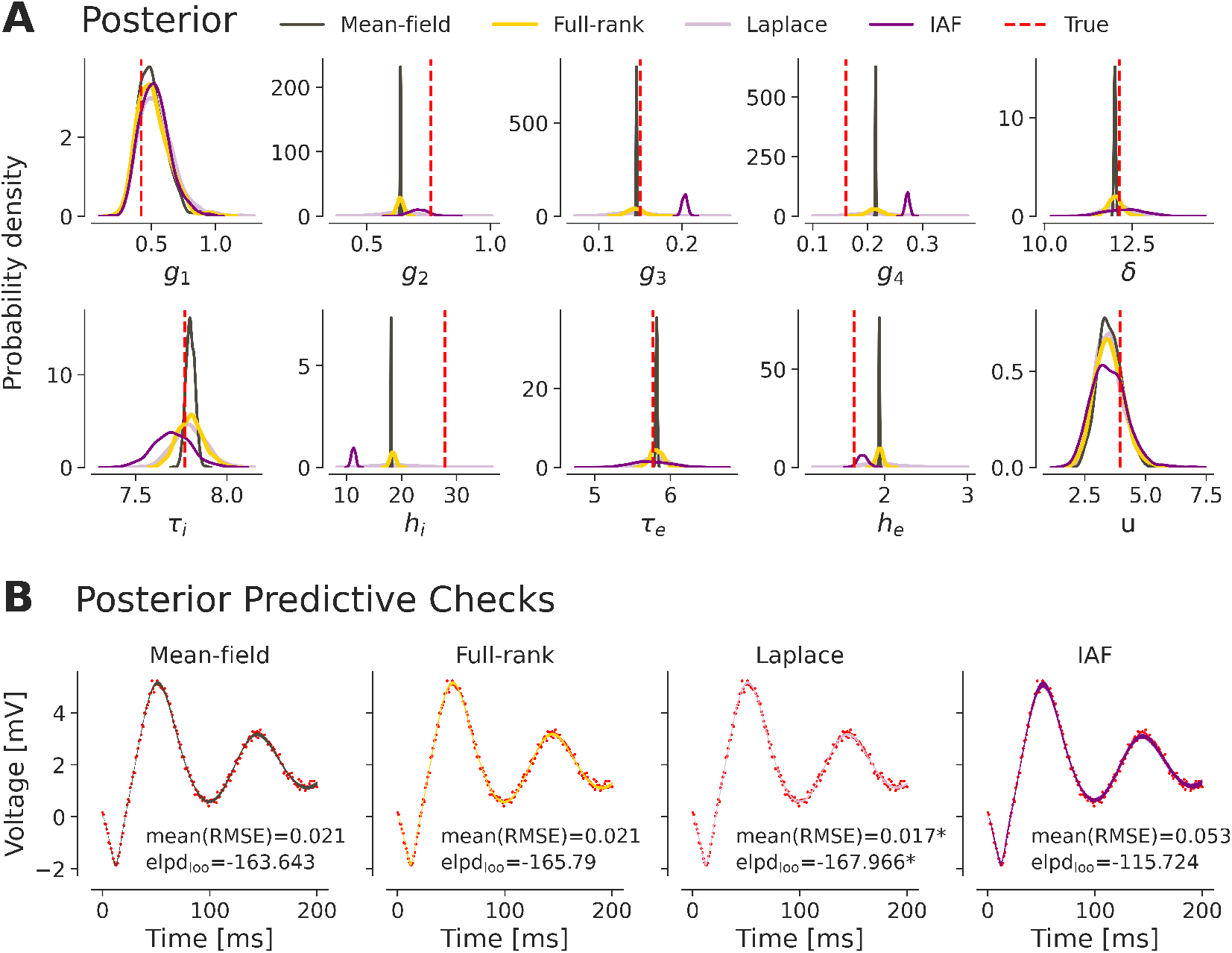
Different modes in the parameter space can generate similar observations. (**A**) Meanfield (in brown) and full-rank (in yellow) variational inference, Laplace approximation (in mauve) and Inverse Autoregressive Flow (IAF) variational inference (in purple, with *k* = 1 flow) target different modes of the parameter space, that produce similar dynamics to the true data generative process (in red). (**B**) Posterior predictive checks are generated from the model using the (1000 samples) randomly drawn from estimated posterior distribution (one line correspond to the dynamics given by one sample from the joint posterior, in which all overlapped). The average RMSE to the data and the Pareto-smoothed importance sampling approximation of the expected log pointwise predictive density (elpd_loo_), which the later approximates cross-validation, are reported for each inference method. The Laplace approximation, which encompasses the true data generative parameters, has, to a very limited extent, the best predictive power.

**Figure S2.**
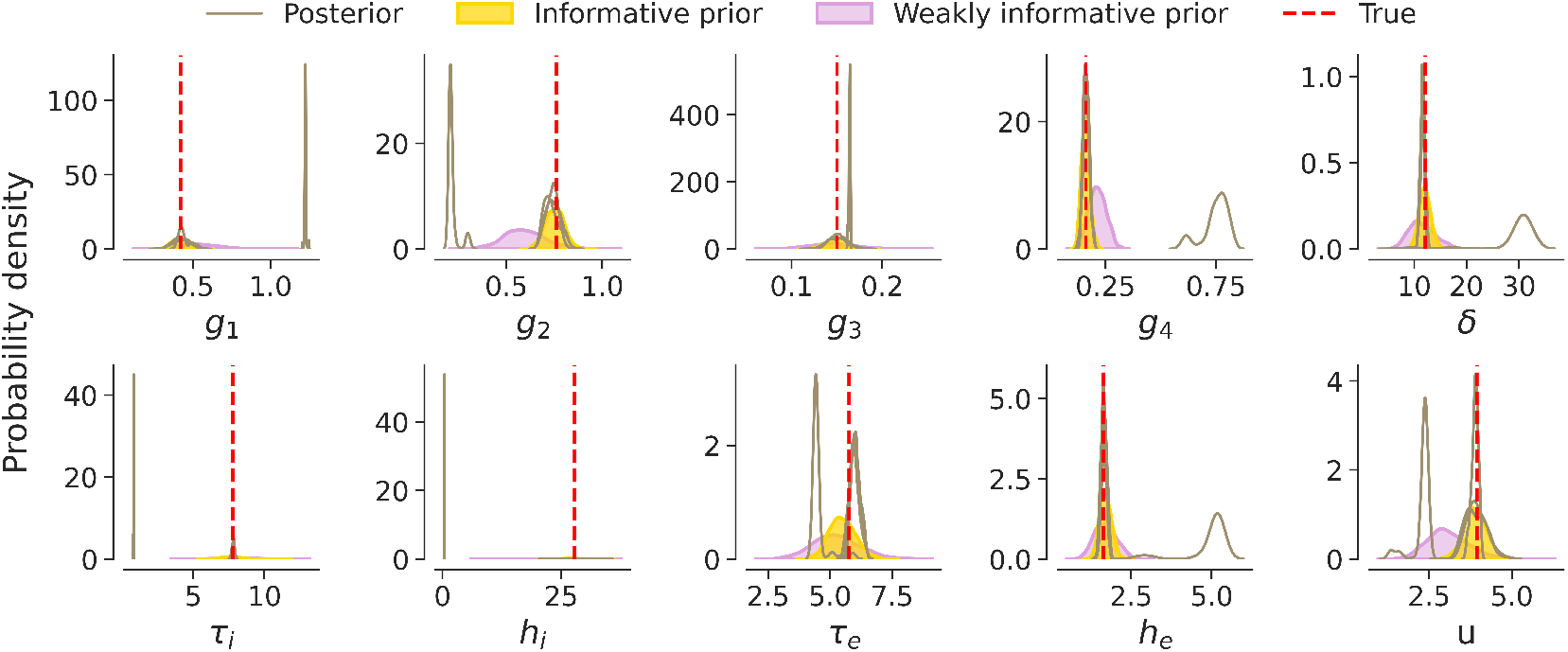
Enforcing more informative priors does not solve multi-modality. Replacing the reference weakly informative priors described in Table 1 (mauve) with informative priors (yellow) described in Table S1, is not sufficient to achieve convergence of inference (here HMC chains, in light brown).

**Figure S3.**
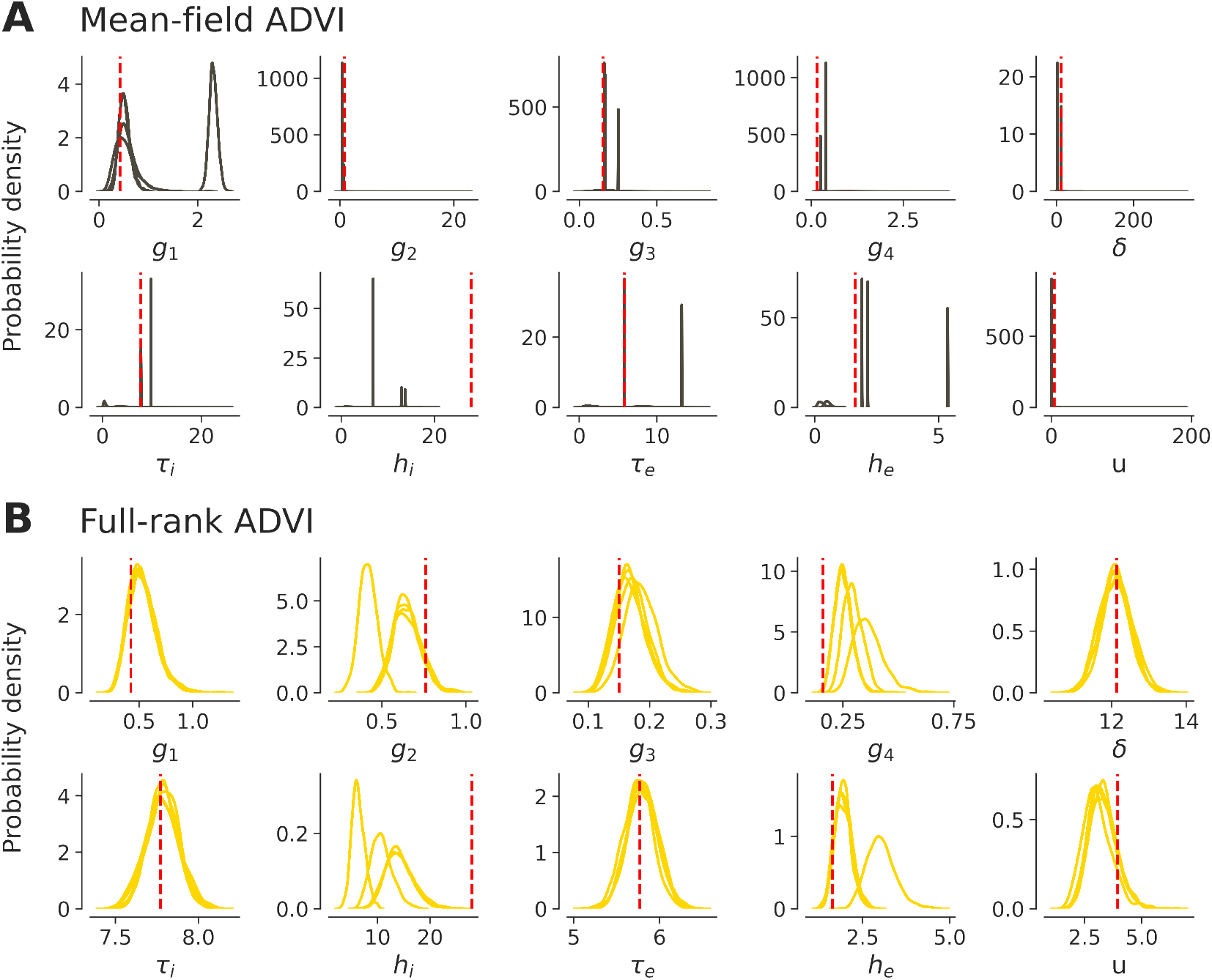
(**A**) Mean-field and (**B**) full-rank variational inference exhibit multi-modality when run multiple times with random initialization. Estimated posterior distributions are plotted as solid lines and the true data generative parameters are shown as red dashed vertical lines.

**Figure S4.**
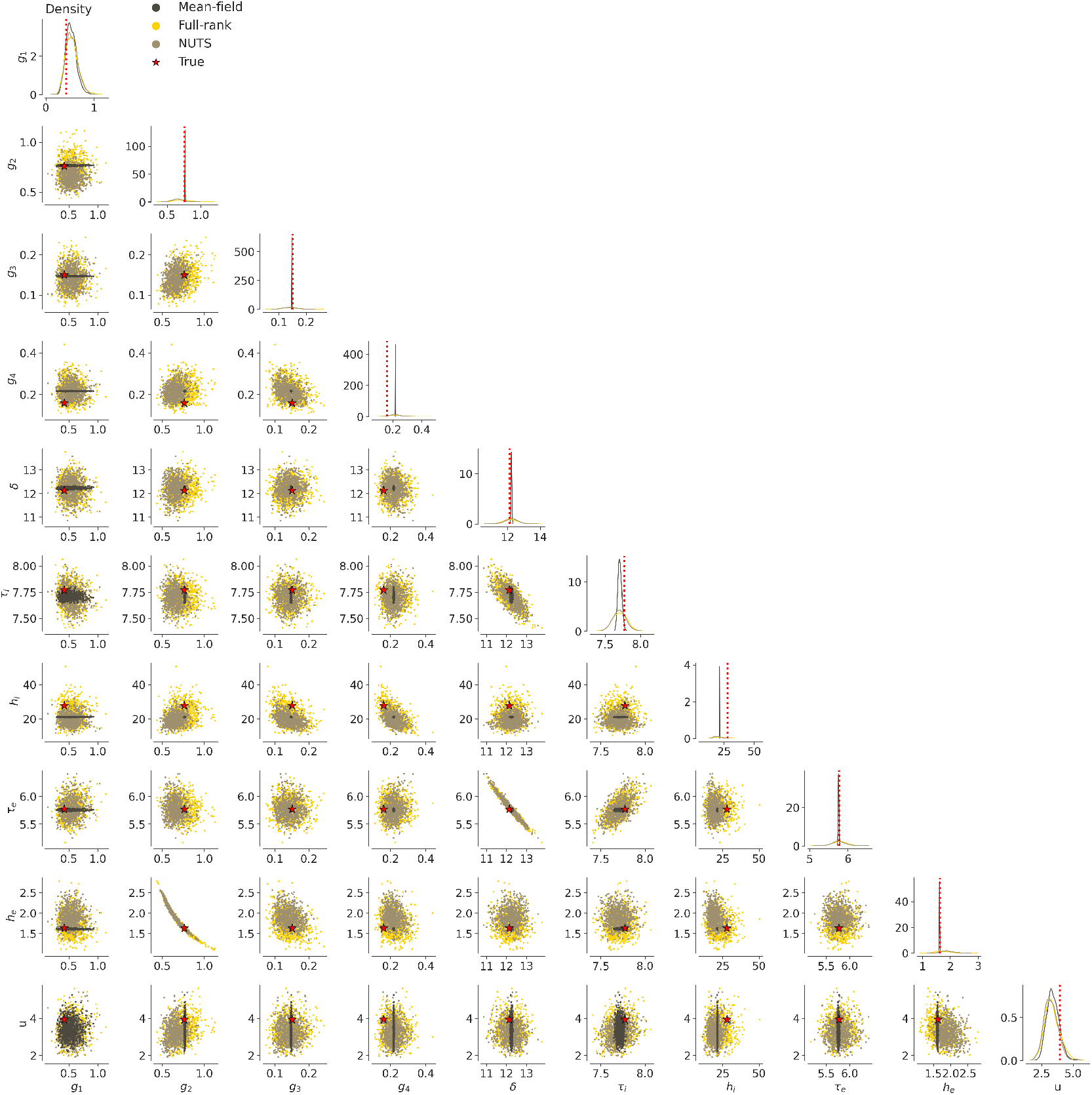
Marginal densities (diagonal) and pairplots (lower triangle) of posterior samples obtained with Stan’s implementation of Variational Inference (mean-field ADVI: in dark brown and full-rank ADVI: in yellow) and HMC (NUTS: in light brown). Full-rank ADVI and NUTS both provide posterior samples that encompass the true generative parameters (in red) and capture complex geometries in the posterior.

**Figure S5.**
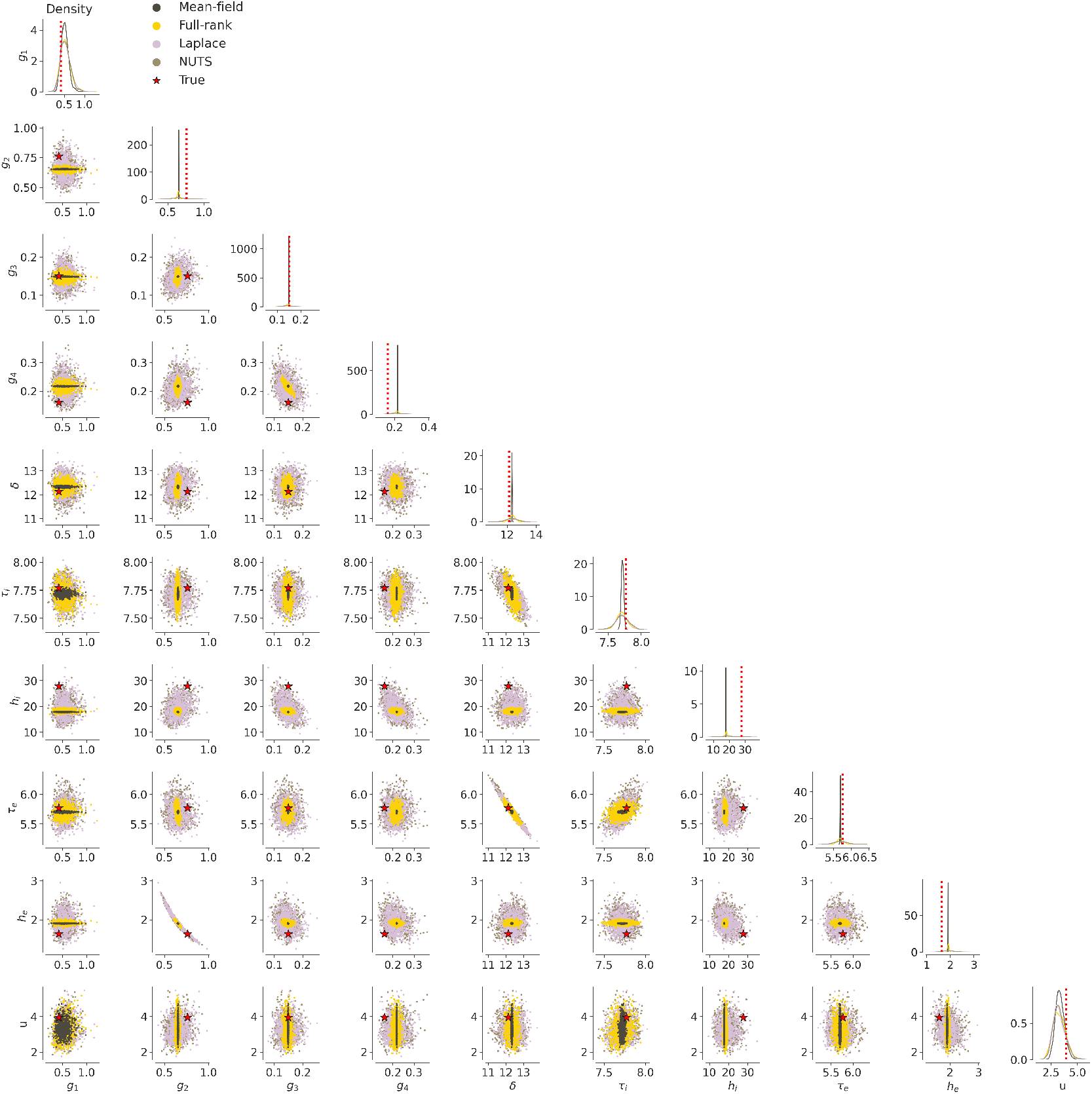
Marginal densities (diagonal) and pairplots (lower triangle) of posterior samples obtained with Numpyro’s implementation of Variational Inference (mean-field ADVI: in dark brown, full-rank ADVI: in yellow, automatic Laplace approximation: in mauve) and HMC (NUTS: in light brown). Laplace approximation and NUTS both provide posterior samples that encompass the true generative parameters (in red) and capture complex geometries in the posterior. For some parameters, full-rank VI provides overconfident posterior distributions which overlook the true parameter value.

**Figure S6.**
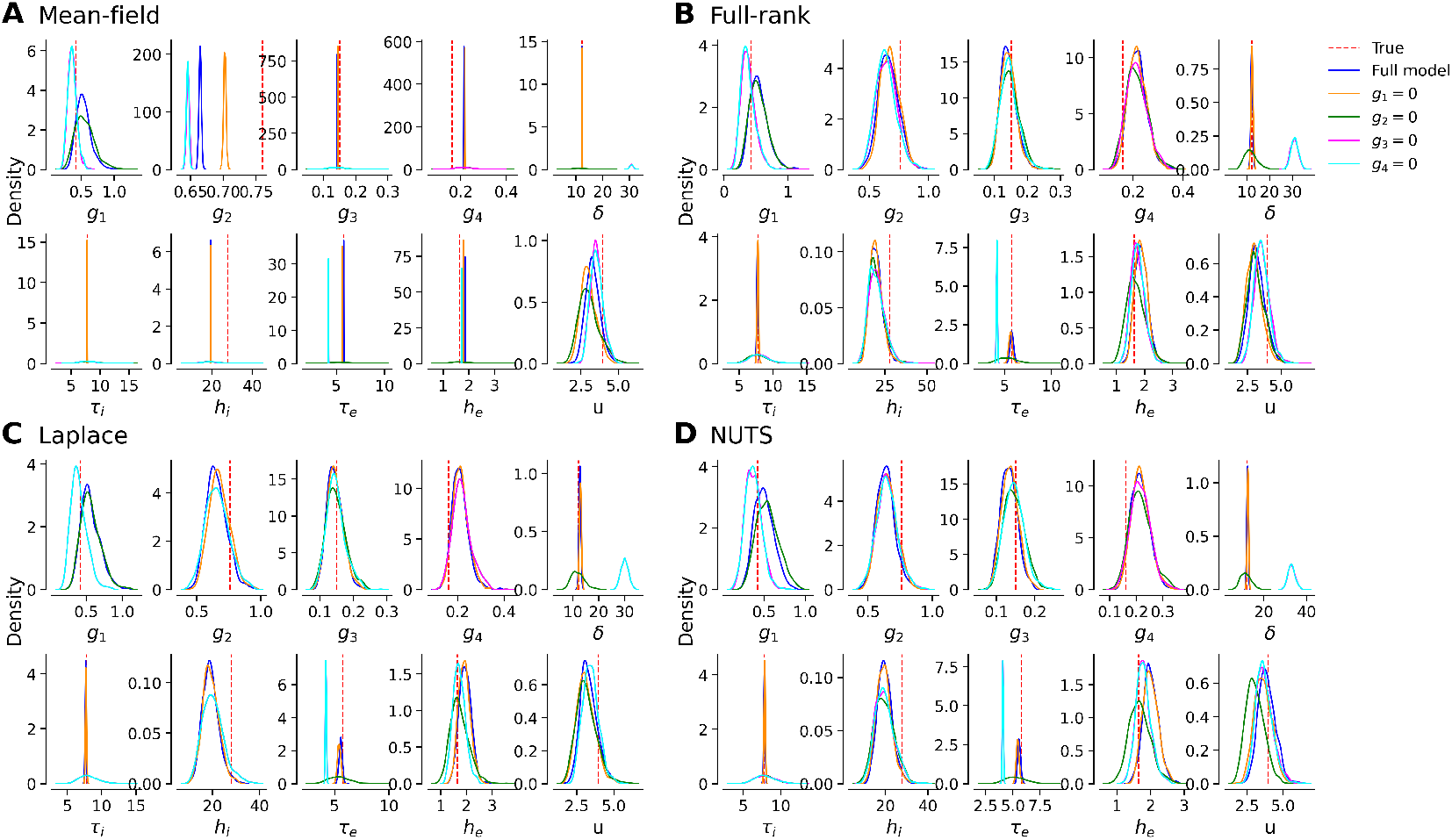
Posterior distributions of parameters from inference on subset models where a single connectivity parameter is set to zero (full model is in blue, true values in red), performed with (**A**) Mean-field ADVI, (**B**) Full-rank ADVI, (**C**) Laplace approximation, and (**D**) NUTS.

**Figure S7.**
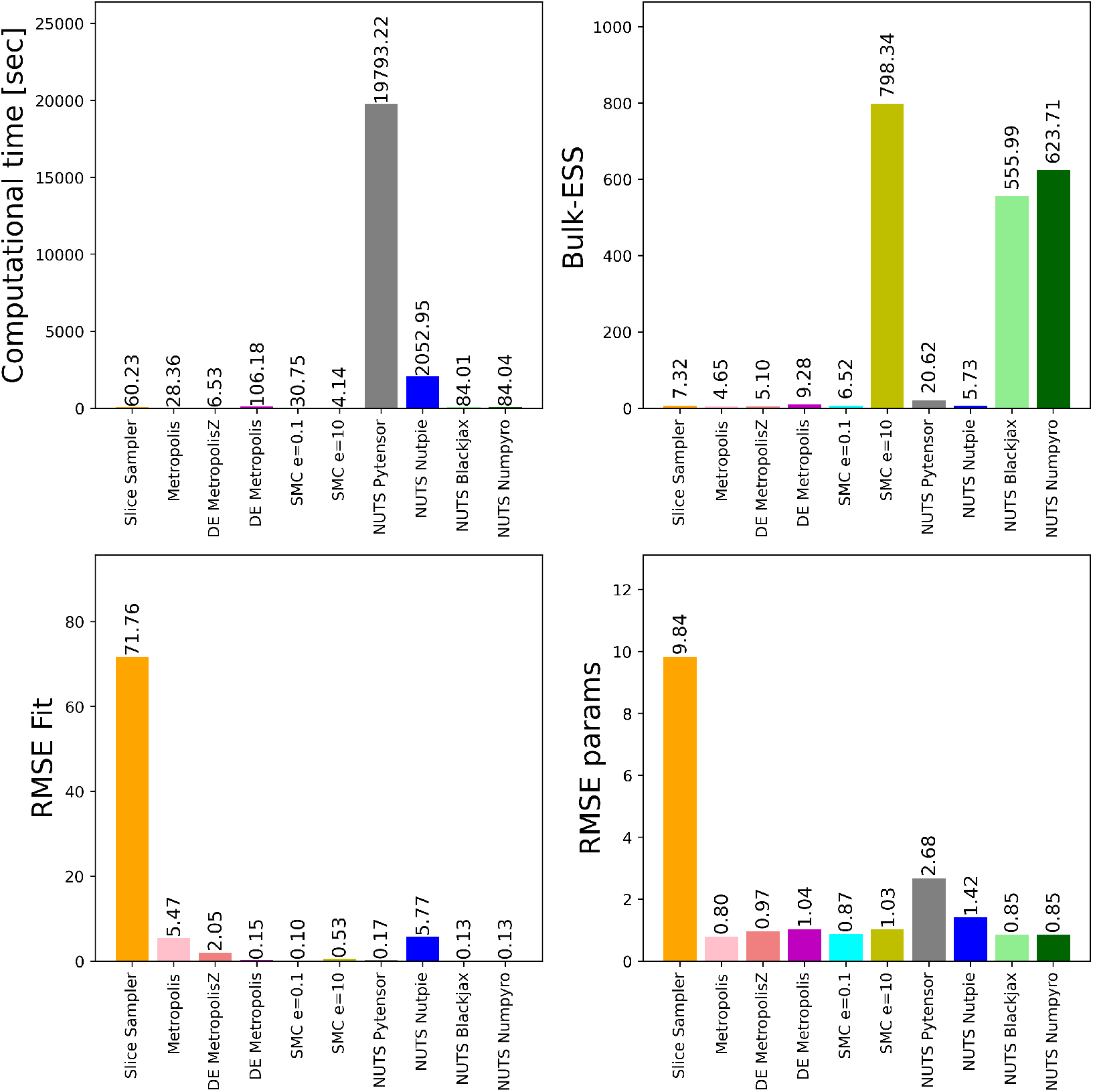
Algorithmic benchmarking in PyMC involved running gradient-free samplers such as Slice sampler, Metropolis, Differential Evolution (DE) Metropolis sampling, Adaptive Differential Evolution Metropolis (DE MetropolisZ), Sequential Monte Carlo (SMC) with different thresholds (e=1, e=10), and NUTS with various backends (Pytensor, Nutpie, Blackjax, and Numpyro). In sum, NUTS (with Blackjax or Numpyro backend) and SMC outperform the Metropolis family algorithms.

**Figure S8.**
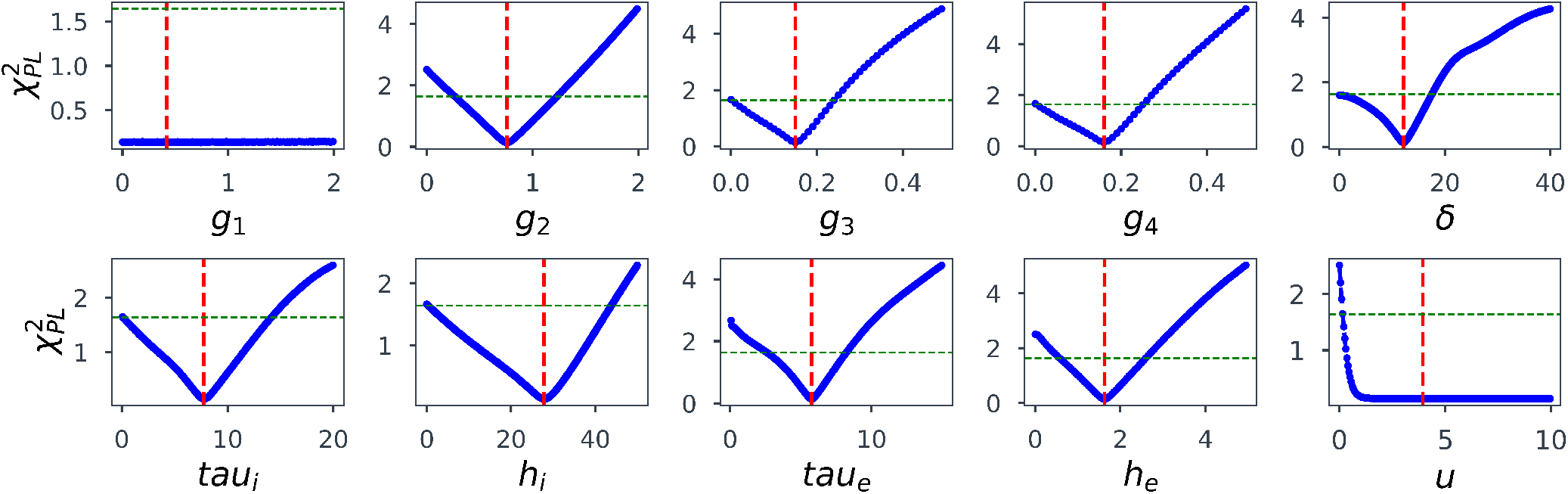
Sensitivity analysis using the profile likelihood (Raue et al., 2011, 2009; Wieland et al., 2021; Hashemi et al., 2023). We calculated the profile likelihood by varying one parameter of interest while fixing all other parameters at their true values (ground truth, dashed red line), and measuring the deviance (e.g., chi-squared error) between the observed data and the model output. This process estimates confidence intervals for the parameter of interest, based on the observed data. According to the profile likelihood, identifiable parameters have finite confidence intervals, structurally non-identifiable parameters show infinite confidence intervals in both directions (such as *g*_1_), and practically non-identifiable demonstrate finite confidence interval in only upper bound (such as *u* parameter). The threshold Δ_*α*_ = *χ*^2^(*α, df*) to determine non-identifiability (dashed green line) is the *α* quantile of the *χ*^2^-distribution with *df* = 1 degrees of freedom, and *df* = #*θ* being the number of parameters, corresponding to the pointwise and simultaneous confidence intervals, respectively.

